# Bio-synthesis of ^15^N-Labeled G-Quadruplexes to Investigate the Structure and Interactions in the Cell Lysate using Nuclear Magnetic Resonance

**DOI:** 10.1101/2025.11.03.685478

**Authors:** Mengbing Zou, Fujia Tian, Wenwen Zheng, Mengzhen Jiang, Shengkai Zhang, Xiaoli Liu, Changxing Ma, Meiping Zhao, Liang Dai, Conggang Li, Shenlin Wang

## Abstract

G-quadruplexes (G4) play key roles in biology, making it critical to understand their structure and ligand-binding behavior in cellular environments for advancing G4-targeted therapeutics. While in-cell nuclear magnetic resonance (NMR) is a powerful technique for studying G4 in situ, its application is limited by the challenge of producing isotope-labeled single-stranded DNA (ssDNA). Here, we introduce Restriction Endonuclease Digestion (RED), a simple and cost-effective method to generate ^15^N-labeled ssDNA. This approach combines molecular cloning and enzymatic design processing by propagating plasmids in *E. coli* cultured with ^15^NH_4_Cl, followed by double restriction digestion and isolation target ^15^N-ssDNA. Using RED, we produced milligram-scale quantities of 96%-enriched ^15^N-labeled human telomeric G4 ssDNA (wtTel23c, CTAGGG(TTAGGG)3), ideal for NMR analysis. The NMR spectra revealed that wtTel23c adopts G4 topology and undergoes multiple conformations of wtTel23c in potassium-containing solutions and in *Xenopus laevis* cell lysate. Interaction studies with the ligand TMPyP4 showed distinct binding profiles in cellular and dilute environments. In dilute solution, TMPyP4 binds to the top tetrad of wtTel23c, while it binds to the loop in cellular environments. The RED method offers an efficient strategy for producing stable isotope-labeled ssDNA, opening new avenues for studying G4 structures and their ligand interactions in complex biological contexts.

## 1. Introduction

G-rich single-stranded DNA (ssDNA) has the ability to adopt G-quadruplexes (G4) in living systems[1,2]. G4s play crucial roles in various physiological processes, e.g., protection against chromosomal degradation, end-to-end fusion, and regulation of gene replication, transcription, and translation [3-5]. As G4s are widely found in the promoter regions of cancer related genes, many G4s are potential drug targets for cancer therapy [6-9]. Designing ligands targeting G4s has been proposed as an effective strategy for drug discovery in this context. Structural biology approaches have made significant progress in elucidating the structures of G4s and complexes with their ligands, providing valuable clues for drug design[10-12]. However, determination on the structures of G4-ligand complexes in the complex cellular environment remains challenging. It is particularly critical for G4 that exhibit polymerization characteristics, such as telomeric G4, which alters its conformations depending on solution conditions and ligand-binding states. Therefore, the structure characterization of G4 in cellular conditions or evaluation the consistence of G4 structures between in diluted solution and in cellular environment is of utmost importance in current G4 research. *In-cell* nuclear magnetic resonance (NMR) has emerged as a powerful tool for characterizing biomolecular structures in their native cellular environment and has been applied to DNA studies [13-20]. Salgado et al. investigated how the intracellular environment modulates the interaction between tetramolecular DNA G4 and ligands [21]. Furthermore, Hansel et al. examined the formation of higher-order G4 structures, consisting of stacked G4 subunits in a *Xenopus laevis* oocyte model[22].

Albeit the successful examples, however, in-cell NMR has not been applied widely for studying G4s interactions with their ligands in cellular environments. One of the main challenges is the difficulty in obtaining uniformly isotope-labeled single stranded DNA (ssDNA), which is necessary for distinguishing the NMR signals of the G4 of interest from those of other vast amount natural abundance cellular components. ^15^N-labeled ssDNA is mainly synthesized by solid-phase chemical synthesis or enzymatic catalysis [23-26]. However, the former method is limited by DNA length, and is prohibitively expensive for obtaining milligram quantities of uniformly ^15^N-labeled DNA. Several enzymatic methods, such as the DNA polymerase fill-in reaction and polymerase chain reaction, have been proposed for preparing labeled DNA [23,27-30]. Nevertheless, these methods also rely on expensive ^13^C and/or ^15^N-labeled deoxynucleoside diphosphates (dNTPs) as precursors. Additionally, synthesizing GC-rich ssDNA or DNA with repeating units poses difficulties due to their potential to block polymerases owing to formation of secondary structures [31]. Production of isotope labeled double stranded DNA can be achieved by propagation of plasmids in *E. coli* cells with subsequent enzymatic digestions.[32] However the isotope labeling on single stranded DNA by bio-synthesis methods are still lacking.

Therefore, there is a demand for new economic and robust bio-synthesis approaches to produce ^15^N-labeled ssDNA. In this study, we present a novel method for the preparing ^15^N-labeled ssDNA based on plasmid propagation in *Escherichia coli* (*E. coli*) for in-cell NMR studies. The strategy involved propagating plasmids in *E. coli* and enzymatic digestion to recover the double-stranded DNA (dsDNA) for ssDNA production. The target DNA sequences were designed to include different restriction enzyme digestion sites at the 5’ and 3’ ends, resulting in asymmetrical dsDNA. Subsequently, the ssDNA was separated and purified from the dsDNA. To obtain ^15^N-labeled ssDNA, the ^15^N-NH_4_Cl was supplemented as sole nitrogen source, resulting in a similar method to the production of ^15^N-labeled proteins through heterologous expression. As ^15^N-NH_4_Cl is an economic precursor, the method has unique advantage in low cost compared to other methods for ^15^N-labeled DNA productions. We applied this method to produce a ssDNA with the sequence of CTAGGG(TTAGGG)_3_, wtTel23c. This sequence adopts the G4 structure and represents the repeats of the short DNA sequence present in the G-overhang architecture of telomeric DNA. A sample of milligram quantities of ^15^N-labeled wtTel23c with 96% ^15^N-labeling enrichment were prepared. We utilized the ^15^N-labeled wtTel23c to investigate its G4 topology and its interactions with the TMPyP4 ligand in both in dilute solution and in the *Xenopus laevis* oocyte extracts. In a K^+^ containing solution, wtTel23c adopted a hybrid-1 G4 topology, and displayed polymorphic conformations that interconverted with each other. The presence of the *Xenopus laevis* cell lysate affects the the proportion distribution of conformational states of wtTel23c. Through NMR and fluorescence spectroscopy, we found that TMPyP4 binding decreased the polymorphism of wtTel23c. The binding sites of TMPyP4 and wtTel23c exhibited significant differences between the in vitro and the cell lysate environments. In the dilute solution, TMPyP4 bound to wtTel23c at the top G-tetrad region, whereas in the cell lysate environment, the interaction occurred via loop attachment. The availability of ^15^N-labeled ssDNA provides valuable opportunities for gaining insights into the binding mechanism of G4 and ligands.

## 2. Methods

### 2.1. Construction of the recombinant pUC57-wtTel23c plasmid

The plasmid containing the human telomeric G4 sequence (pUC57-wtTel23) was ordered from GenScript (Nanjing, China). The pUC57-wtTel23 plasmid was constructed by ligating the desired sequence (1032 bp) into the BsmBI restriction enzyme sites of the pUC57 vector (2.6 kb) (Fig 1). The 1032-bp DNA sequence contained 12 repeats of the TAGGG(TTAGGG)_3_ (wtTel23) sequence with short linker sequences between each repeated unit. The KpnI and BamHI recognition sites were present at the 5′ and 3′ ends of wtTel23, respectively. To generate asymmetrical double-stranded DNA (dsDNA), a double enzymatic digestion was performed. The ^15^N-labeled ssDNA was prepared using DH5α cells harboring the target plasmid, following the procedure described below.

**Figure 1.**
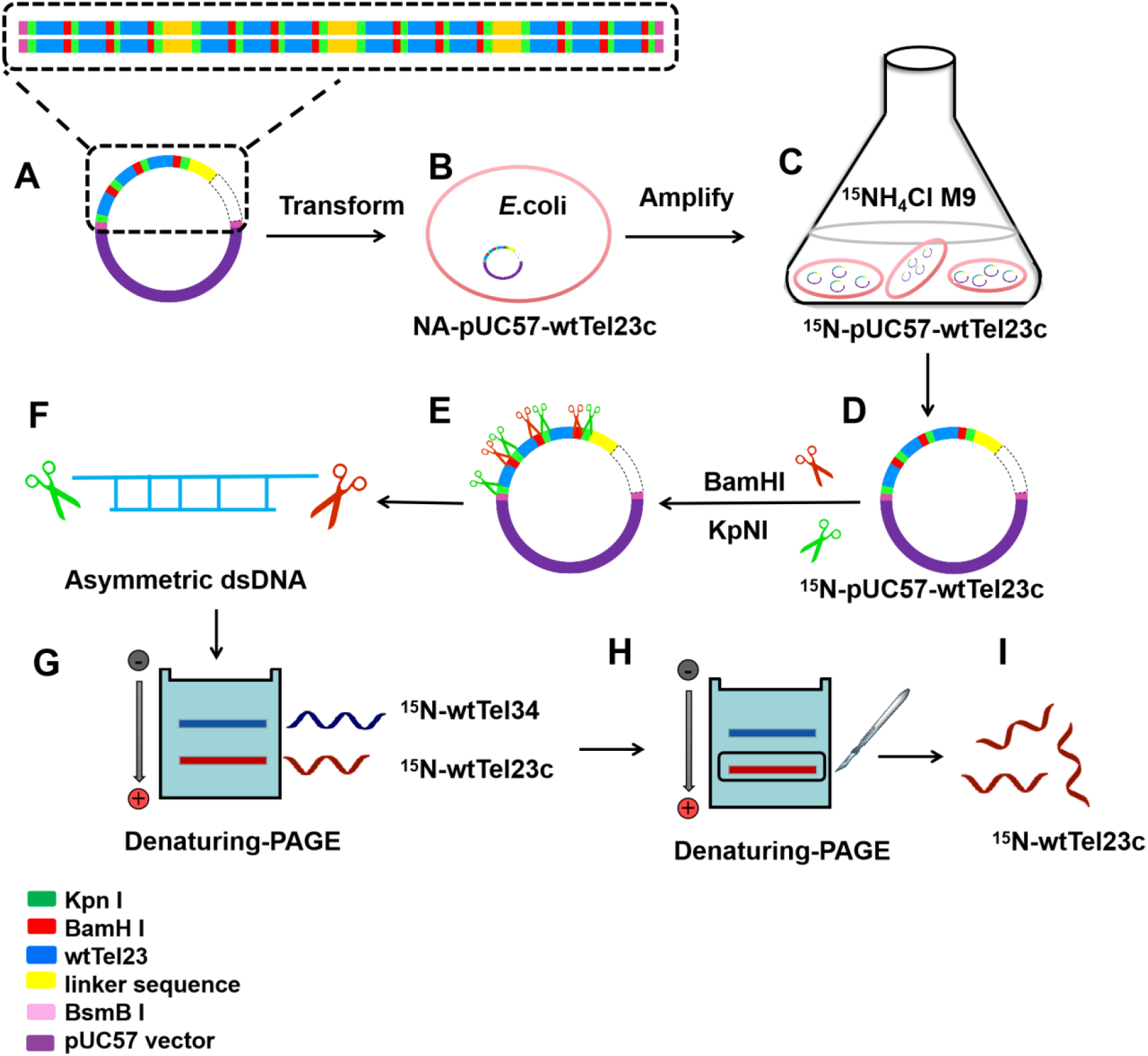
The schematic flow of RED-ssDNA biosynthesis. (A) Overall design of the plasmid for replication by *in vivo* cloning to obtain ^15^N-labeled ssDNA. The target DNA fragments of wtTel23 are shown in blue rectangle. The restriction enzyme sites of Kpn I (green) and BamH I (red) are at the 5′ and 3′ ends, respectively. Fifteen repeats of the target genes are connected to the vector in series, and every three repeats are connected with a linker with random sequence (yellow) to facilitate the plasmid construction. The tandem of all DNA fragments are ligated into the pUC57 vector at BsmB I restriction enzyme sites to form the plasmid pUC57-wtTel23c. (B) The constructed plasmid is transformed into DH5α *E. coli* cells for propagation. (C) The transformed DH5α *E. coli* cells with pUC57-wtTel23c are grown in M9 medium with ^15^N-labeled NH_4_Cl as the sole nitrogen source to produce the ^15^N-labeled pUC57-wtTel23c plasmid. (D, E) The purified ^15^N-labeled pUC57-wtTel23c plasmids are treated with the restriction endonucleases Kpn I and BamH I. (F) The digested products are obtained as asymmetrical dsDNA. (G) The asymmetrical dsDNA are further separated and purified by urea-PAGE (45 cm × 35 cm). (H) The target ssDNA are recovered and electroeluted to obtain the target ^15^N-labeled ssDNA.

### 2.2. Preparation of ^15^N-labeled ssDNA wtTel23c

Preparation of ^15^N-labeled ssDNA wtTel23c was carried out using the biosynthesis method. Clones of DH5α *E*.*coli* cells harboring the pUC57-wtTel23c plasmid were cultured at 37°C in 50 mL LB medium supplemented with 100 mg/mL ampicillin. After 5 hours of growth, the cells were transferred to M9 minimal medium containing 1.0 g/L ^15^N-labeled NH_4_Cl as the sole nitrogen source. When the OD_600_ reached 1.0, chloramphenicol was added to a final concentration of 18 mg/mL. The cells were harvested 9 hours after medium exchange. Plasmids were extracted using a previously reported procedure[33]. The yield of extracted plasmid was approximately 10 mg/L of cell culture. The extracted plasmids were digested with BamH I and Kpn I enzymes to obtain asymmetrical dsDNA. Following the enzyme digestion reaction, 6% polyethylene glycol 6000 was added to remove the DNA of the pUC57 vector. The supernatants containing the asymmetrical dsDNA was denatured and separated by 12% denaturing acrylamide gel electrophoresis (urea-PAGE). The two ^15^N-labeled ssDNAs, referred as wtTel23c [5′-CTAGGG(TTAGGG)_3_-3′] and wtTel34 [5′-GTAC(CCCTAA)_3_CCCTAGGTAC-3’], were collected and recovered from the gel by incubation with deionized water. The collected wtTel23c was changed buffer to G4 folding buffer (25 mM potassium phosphate, 75 mM KCl, pH = 7.0).

### 2.3. Circular dichroism (CD) spectroscopy

CD spectra were recorded using a Jasco J-810 spectrometer. The molar ellipticity, [θ] (degrees cm^2^ dmol^−1^), was calculated using the following equation: [θ] = 10^6^ × [θ]obs × C^−1^ × L^−1^, where [θ]obs represents the ellipticity (mdge), C is the molar concentration of oligonucleotide, and L is the optical path length of the cell (cm). The spectra were collected at a temperature of 25°C using a cell with a path length of 0.1 cm. Each sample had a DNA concentration of 20 μM. Spectra were acquired with a bandwidth of 2 nm, a scan rate of 50 nm min^−1^, and a step size of 1 nm, with a 2-second measurement time per point. Signal-averaged spectra were obtained by scanning each sample at three times. Baseline correction was performed by subtracting the spectra of the buffer. To obtain the spectra of the DNA–TMPyP4 complex, the DNA samples were supplemented with TMPyP4 at a 1:1 molar ratio and incubated at 4°C overnight, before measuring the CD spectra.

### 2.4. Cell lysate sample preparation

Cytoplasmic extracts of *X. laevis* oocytes were prepared as previously described[22]. Briefly, *X. laevis* oocytes were pelleted by centrifugation at 400 × *g* for 1 minute, and the supernatant was removed. The oocytes were then centrifuged at 12,000 × *g* for 5 minutes. To obtain a homogenized extract solution, the oocytes were crushed on ice using a glass pipette. Finally, the homogenized oocytes were centrifuged at 12,000 × *g* for 30 minutes to collect the crude extract, which was subsequently freeze-dried into powder form.

### 2.5. Steady-state fluorescence spectroscopy

The interaction between TMPyP4 to wtTel23c was investigated using steady-state fluorescence spectroscopy with 2-aminopurine as the probe. Fluorometric experiments were conducted at 25°C using the Jasco FP-7000 spectrofluorometer. The ssDNA containing 2-aminopurine modifications were excited at 305 nm (with a bandwidth of 5 nm), and the emission spectra (bandwidth of 5 nm) were measured from 320 to 420 nm at 1-nm intervals. Oligodeoxynucleotides modified with 2-aminopurine (at a concentration of 400 μM) were utilized to study the binding between TMPyP4 and wtTel23c in diluted solution as well as within cell lysate environment. The same concentration of cell lysate was used as a control.

### 2.6. NMR spectroscopy

NMR experiments were conducted using Bruker 700-MHz, 800-MHz, 900-MHz and/or 950-MHz spectrometers equipped with a cryogenic triple resonance probe. The G4 samples were dissolved and folded in G4-folding buffer at pH 7.0 (75 mM KCl, 25 mM potassium phosphate, and 10% D_2_O). To ensure proper folding, G4 was annealed at 95°C for 10 minutes and chilled on ice. Both *in-vitro* and *in-cell* samples were loaded into 5-mm Shigemi tubes. I*n-vivo* experiments were carried out at approximately 22°C to minimize any sample alterations. To study the G4– TMPyP4 samples, the G4 was mixed with TMPyP4 solution at a final molar ratio of 1:1. The mixture was then incubated at 37°C for 2 hours before measurement. Two-dimensional (2D) ^1^H-^15^N band-selective optimized flip-angle short-transient heteronuclear multiple quantum coherence (sfHMQC) spectra were collected for both ^15^N-labeled and natural abundance G4. Selective ^1^H excitation pulses were centered at 11.2 ppm, and the inter-scan delay was set as 200 ms. The concentrations of the DNA samples used for the *in-vitro* study ranged from 0.1 to 1.5 mM.

### 2.7. Modeling of TMPyP4-wtTel23c complex

We constructed the ligand free and the TMPyP4 bound form of wtTel23c using the NMR structure of DNA G4 with a mixed hybrid-type geometry as the starting template (PDB ID: 2HY9 from the Protein Data Bank) [34]. The structure was modified to obtain a G4 structure with the sequence 5’-CTAGGGTTAGGGTTAGGGTTAGGG-3’. The ligand free wtTel23c was then solvated in a truncated rectangle water box where the solute spread 10 Å apart to the boundary of box. The ligand molecule was placed as least 10 Å away from the DNA to ensure there are three layer of water molecules between DNA and the ligand. The solute was finally neutralized by counterion, K^+^, leading to desired K^+^ concentration with more adding cations. The AMBER OL15 [35] force field was employed to represent the DNA structure, water molecules were described by TIP3P model and K^+^ ions was represented by the Joung-Cheatham model [36]. The partial charges of TMPyP4 were obtained from the restrained electrostatic potential (RESP) method after obtaining the electrostatic potential using Gaussian 09, and the force field parameters of TMPyP4 were generated from the AMBER GAFF2 [37] force field.

### 2.8. Molecular dynamics simulations

All simulations were performed using the GROMACS 2021 software package. The simulation system first underwent the energy minimization in two steps: (i) minimization of water and counterions with restraints (1000 kJ/mol/nm) on DNA and ligand; (ii) minimization of the whole system without restraints. After energy minimization, the systems was subjected to three equilibration steps: (i) a 100 ps run in the canonical (NVT) ensemble to heat from 0 K to 295 K with restraints on DNA and ligand; (ii) a 1 ns run in the isobaric-isothermal (NPT) ensemble with decreasing restraints (1000, 800, 600, 400 and 200 kJ/mol/nm) on DNA and ligand; (iii) a 10 ns run in the NPT ensemble without restraints. Finally, an unrestrained production run at 295 K was performed in the NPT ensemble for 1 μs.

We used the LINCS [38] algorithm to constrain all bonds involving hydrogen atoms. Periodic boundary conditions were used in all three dimensions. A 1.0 nm cut-off was applied for short-range nonbonded interactions. The Particle-Mesh Ewald (PME) [39]method was used for long-range electrostatic interactions. The temperature of simulation system was controlled using the V-rescale [40] thermostat with a relaxation time τ = 0.1 ps and the pressure was kept at 1 atm using the Parrinello-Rahman pressure-coupling with a relaxation time τ = 2.0 ps and compressibility 4.5×10^−5^ bar^-1^. The time step is 2 fs and the coordinates of simulation system were saved every 5000 steps, enabling a 10 ps interval for analysis.

## 3. Results and discussion

### 3.1. Plasmid design to produce ^15^N-labeled wtTel23c

Milligram quantities of uniformly ^15^N-labeled G4 is a prerequisite for NMR-based in vivo and ex vivo studies on G4. We developed a method called RED-ssDNA (Restriction Endonuclease Digestion-ssDNA) biosynthesis method to produce uniformly ^15^N-labeled ssDNA, which relied on *E. coli* propagation. The schematic representation of the RED-ssDNA biosynthesis is depicted in Fig 1. The target DNA sequences containing restricted enzymatic sites were cloned into the pUC57 plasmid and transformed into *E. coli* cells. To produce the ^15^N-labeled plasmids (pUC57-wtTel23c), M9 medium supplemented with ^15^N-NH_4_Cl as the sole nitrogen source was used. Enzyme digestion of the corresponding dsDNA sequences generated asymmetrical dsDNA, from which the desired ssDNA could be obtained by separation using urea-PAGE. To enhance the yield of the target DNA, multiple copies of the target sequences were sub-cloned into the plasmid.

This approach was tested to produce ^15^N-labeled human telomeric G4 with target sequence of wtTel23 [5′-TAGGG(TTAGGG)_3_-3′] that was incorporated into the pUC57 plasmid. Twelve copies of sequences with fifteen tandem wtTel23 repeat units and linker sequences were ligated into the pUC57 plasmid with BsmBI restriction enzymatic digestion sites placed at the 5′- and 3′-ends of the entire sequence (Table S1). The recombinant plasmid pUC57-wtTel23 was then transformed into DH5α *E. coli* cells for propagation in ^15^N-M9 medium. The obtained recombinant plasmid pUC57-wtTel23 was sequenced to confirm the fidelity of the experimental design. Agarose gel electrophoresis was also used to verify the size of the connecting clip (Fig S1). The 5′ and 3′-end of the target G4 sequence contain BamHI and KpnI restriction sites, respectively, which were expected to yield an asymmetrical dsDNA with a sequence of [5′-CTAGGG(TTAGGG)_3_-3′], and its complement sequence was [5′-GATC(CCCTAA)_3_CCC′TAGGTAC-3′]. The resulting asymmetrical dsDNA after BamHI and KpnI double enzymatic digestions was examined by 12% native PAGE, yielding a molecular weight in agreement with the expectation (Fig S1). Subsequently, the asymmetrical dsDNA was separated and purified by 12% denaturing urea-PAGE to obtain two ssDNAs. The two ssDNA chains with unequal length migrated distinguishably on denaturing urea-PAGE, allowing for their separation.

### 3.2. Characterization of ^15^N-labeled ssDNA

Agarose gel, liquid chromatography mass spectroscopy (LC-MS), circular dichroism (CD) spectroscopy, and ^1^H-NMR spectra (Fig. 2A-B, S1 and S4) were collected to validate the bio-synthesized wtTel23c. In the presence of 150 mM K^+^, the CD spectra displayed characteristic patterns for the hybrid-1 G4 structure with positive peaks at 285 nm and 265 nm. The NMR spectra of the wtTel23c sequence from RED-ssDNA synthesis and solid-phase synthesis had similar spectra patterns in the imino-region with ^1^H-chemical shifts of 11-12 ppm. The similar CD and ^1^H-NMR spectra of wtTel23c obtained through RED-ssDNA methods and solid-phase synthesis demonstrated that the RED-ssDNA effectively produced wtTel23c with the desired sequence in solution.

**Figure 2.**
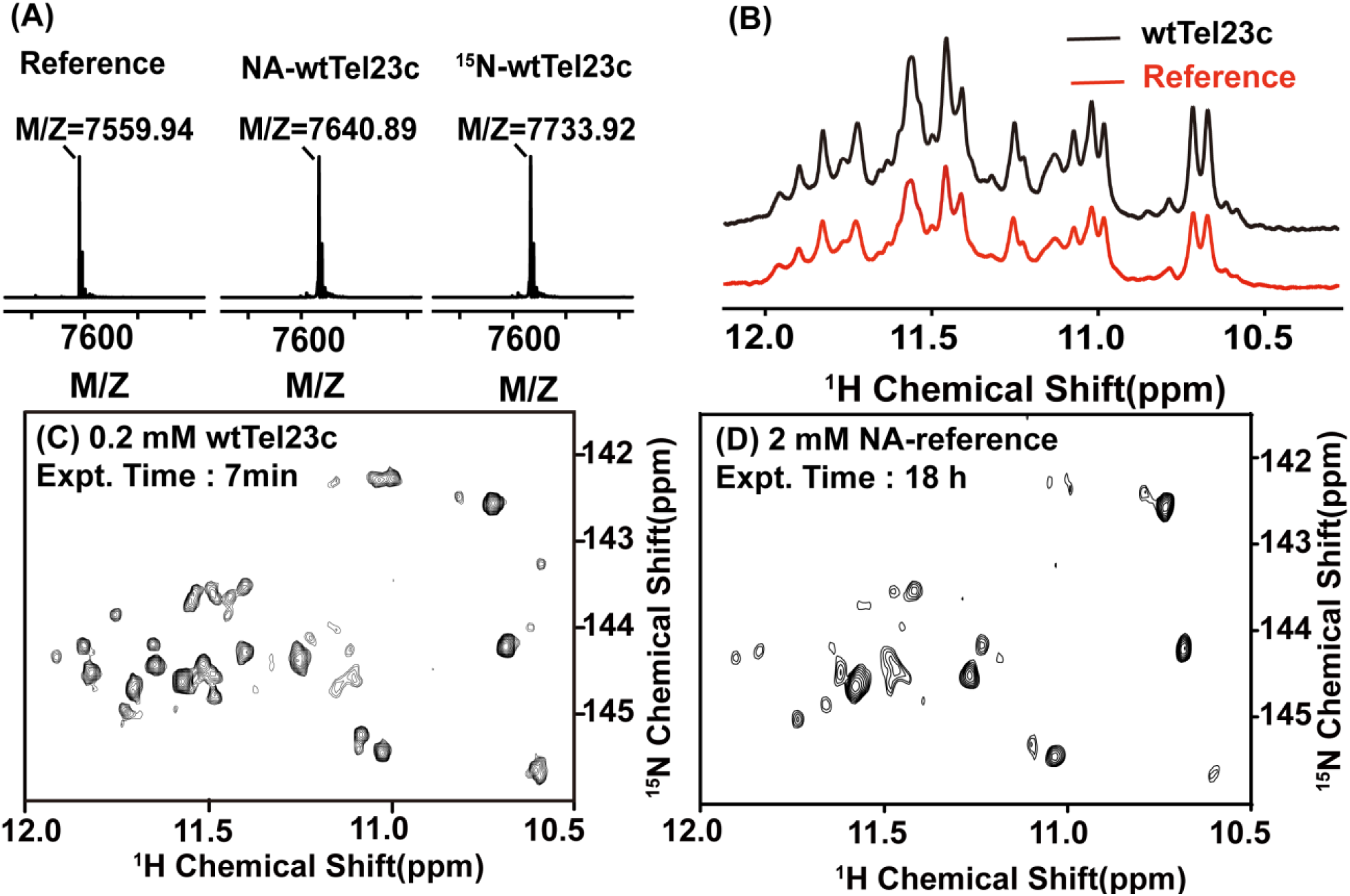
Biophysical characterization of ^15^N-labeled ssDNA. (A) MS analysis of the ^15^N labeling enrichment level of ^15^N-labeled wtTel23c. The reference is wtTel23c synthesized by solid-phase synthesis. Naturally abundant wtTel23c (NA-wtTel23c) and ^15^N-labeled wtTel23c (^15^N-labeled wtTel23c) are prepared by RED-ssDNA biosynthesis. (B) Imino proton regions of 1D ^1^H-NMR spectra of wtTel23c (black) and reference (red), respectively. (C, D) Imino regions of 2D ^1^H-^15^N sfHMQC NMR spectra of ^15^N-labeled wtTel23c and reference, respectively.

LC-MS was used to measure the ^15^N labeling enrichment level of wtTel23c (Fig. 2A). The molecular weight of the natural abundance (NA) wtTel23c and the ^15^N-wtTel23c, both produced using RED-ssDNA methods, were 7640.89 and 7733.92 Da, respectively, showing that ^15^N-wtTel23c had a 93.92 Da higher molecular weight than NA-wtTel23c, corresponding to 96% ^15^N enrichment. MS data for wtTel23c from solid-phase synthesis yielding a molecular weight of 7559.94 Da (Fig. 3D–E), which was a discrepancy of 80.95 Da between wtTel23c produced by solid-phase synthesis and the RED-ssDNA method. It may be attributed to the presence of a phosphate group at the 5’ terminus of wtTel23c by enzymatic digestion, while the 5’ terminus of wtTel23c produced by solid-phase synthesis contained a hydroxyl group.

**Figure 3.**
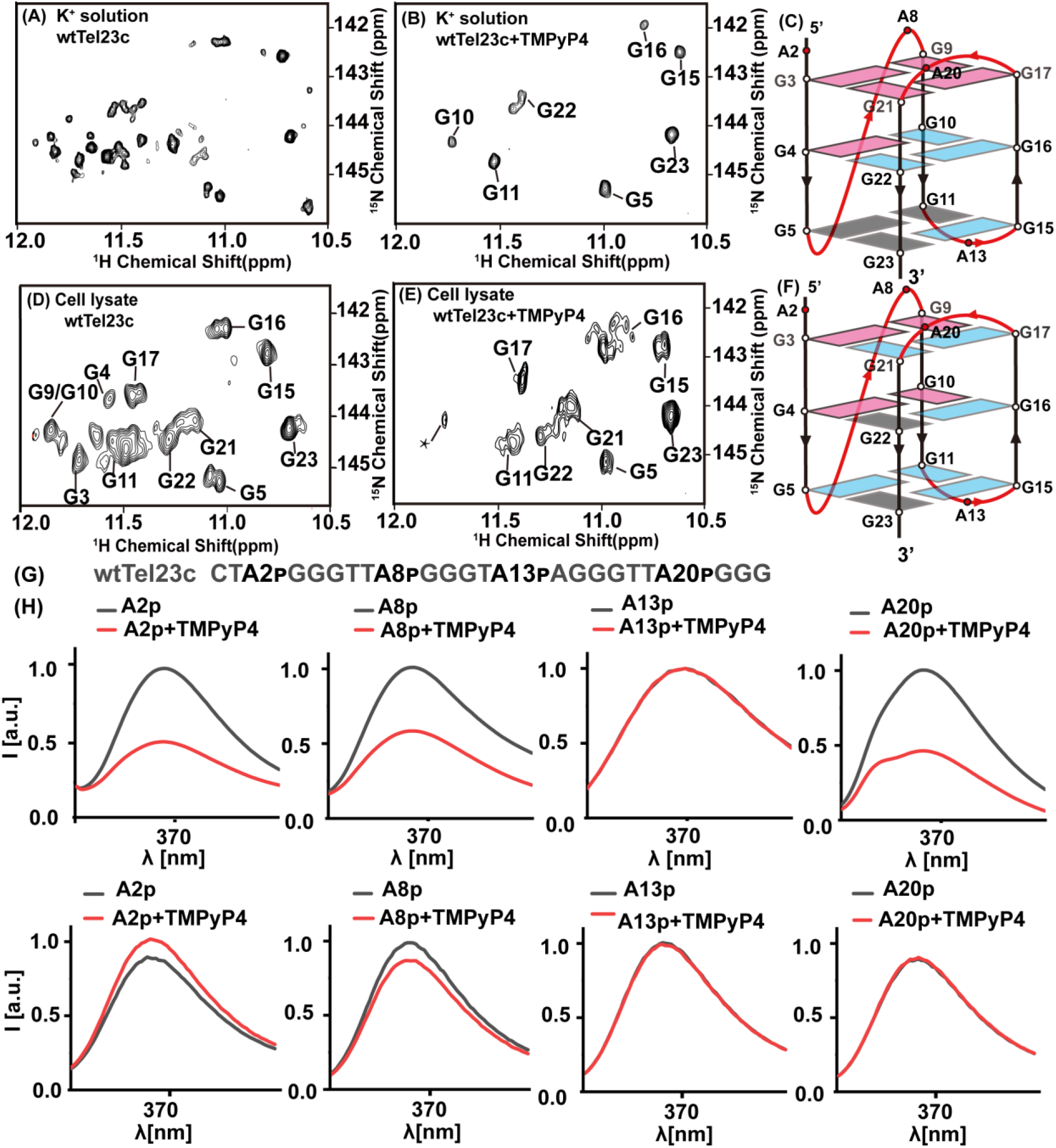
Characterization of the TMPyP4-binding region on wtTel23c in dilute solution and in cell lysate. (A- B) 2D ^1^H-^15^N sfHMQC spectrum of wtTel23c (A) and wtTel23c-TMPyP4 (B), both in solution containing 150 mM K^+^. (D-E) 2D ^1^H-^15^N sfHMQC spectrum of wtTel23c (D) and wtTel23c-TMPyP4 (E), both in cell lysate. The assignments are highlighted. The star indicates the unassigned signal, which may be corresponding to G3, G4, G9, G10, belonging to a group of signals that disappear or weaken. (C and F) The cartoon representation shows the guanines in wtTel23c with significant changes after binding with TMPyP4 in K^+^ containing solution (C) and in cell lysate (F). The pink boxes show the signals that are severely weakening or disappearance, the light blue boxes present the guanine signals that undergo peak shifting, and the gray boxes indicate the cross-peaks without any changes. (H) Overlay of steady-state fluorescence emission spectra of free wtTel23c (black) and wtTel23c-TMPyP4 (red), with oligodeoxynucleotides modified with 2-aminopurine at positions A2, A8, A13, and A20 in dilute solution (G) and in cell lysate (H). The 2-aminopurine sites are shown on the top of (G).

A ^1^H-^15^N sfHMQC spectrum was acquired for the ^15^N-labeled wtTel23c at a concentration of 0.2 mM using a 700 MHz NMR spectrometer. High-quality NMR spectra were obtained within 7 minutes of machine time with a signal-to-noise ratio (S/N) of 133 (averaged for the detected signals) (Fig. 2C). Comparably, an S/N of 27 were obtained on the NA-wtTel23c, which took 18 hours for 2 mM sample (Fig. 2D). Thus, ^15^N labeling of wtTel23c achieved a 37,000-fold increase in S/N unit than the NA-wtTel23c, significantly improved the efficiency of acquisition on ^1^H-^15^N correlation spectra, i.e. sfHMQC and HSQC, This advancement enables possibility to characterize G4 topology and interaction within complex cellular environments.

### 3.3. TMPyP4 binding to the top tetrad of wtTel23c in solution

TMPyP4 binding to wtTel23c in dilute solution was investigated using ^15^N-labeled wtTel23c. The ^1^H-^15^N sfHMQC spectra showed more than the expected 12 cross-peaks corresponding to the imino of guanine (Fig. 2A), suggesting that wtTel23c adopts multiple conformations, which is consistent with previous studies showing that the human telomeric G4 topology is influenced by the 5’ flanking sequence (41). The conformation polymorphisms are temperature-dependent; some peaks weakened or disappeared with increasing temperature, while some peak intensities increased (Fig. S2), indicating that the G4 conformations of wtTel23c are exchangeable under ambient temperature conditions. Upon addition of TMPyP4 at a 1:1 molar ratio, the number of peaks decreased to 7 (Fig. 3), indicating that TMPyP4 binds to G4 and reduced the level of polymorphism of the G4 topology or alters the dynamics of conformational equilibrium. Increase in the molar ratio of TMPyP4 with wtTel23c to 1:2 led to severely broadening of the cross peaks, suggesting the presence of multiple binding sites of TMPyP4 on wtTel23c (Fig. S3A). We used 1:1 molar ratio of TMPyP4 and wtTel23c for further NMR studies. Correspondingly, CD spectra of wtTel23c-TMPyP4 complex showed a decrease in molar ellipticity at 265 nm compared to the free state wtTel23c, indicating a decrease in the parallel conformation as reported before [41], while the dominant conformation still being hybrid-1 (Fig S3).

The structure of wtTel23 (PDB code 2HY9) has been previously determined. The sfHMQC spectra of wtTel23c-TMPyP4 showed similar spectral pattern to wtTel23-TMPyP4 (Fig. S4B). Thus, the NMR assignments of wtTel23c were conducted using assignments of wtTel23 as starting point, using the 2D NOESY spectra to obtain sequential connections and guanine assignments (Fig. S3 and S4). In the 2D sfHMQC spectra of wtTel23c-TMPyP4, assignments were obtained for imino groups of seven guanines (G10, G16, G22, G21, G15, G5, and G23) belonging to the middle and bottom tetrad, while the guanines of the top tetrad (G3, G9, G17, and G21) were not detected (Fig. 3C). This suggested that TMPyP4 might stack onto the top tetrad of wtTel23c, in which the binding may disturb the inter-guanine hydrogen bond network of the top tetrad, thus inducing broadening of the corresponding cross-peaks.

Steady-state fluorescence spectroscopy was performed to confirm the TMPyP4 binding sites on wtTel23c. Previous studies have shown that close contact or structural rearrangement upon TMPyP4 binding can change the max emission wavelength of 2-aminopurine at 370 nm[22]. It thus allows the identification of the binding positions of G4 with TMPyP4. Profiles of four wtTel23c labeled with 2-aminopurine at the A2, A8, A13, and A20 positions (termed A2p, A8p, A13p, and A20p, respectively) were obtained (Fig. 3G). The fluorescent signal of 2-aminopurine modified A14 (A14p) is quenched by a nearby guanine and cannot serve as a reliable reference. Therefore, a T13 to A13p mutation was introduced while maintaining the topology of the G4 (Fig. S5). The addition of TMPyP4 resulted in significant quenching of the fluorescence emission from A2p, A8p, and A20p, all of which are near the top tetrad of the wtTel23c. In contrast, the fluorescence emission profile of A13p remained unchanged (Fig. 3H). This result showed that TMPyP4 binds in proximity to the top tetrad, consistent with the conclusion from the NMR data.

### 3.4. Evaluation of the interaction between the wtTel23c and TMPyP4 in cell lysate

To evaluate whether the TMPyP4 binds wtTel23c at the same site in cellular conditions, the ^15^N-labeled wtTel23c was further microinjected into *X. laevis* oocytes. However, high-resolution spectra were not obtained due to the intracellular factors such as viscosity or crowding. Thus, ^15^N-labeled wtTel23c was added in *X. laevis* oocytes cell lysate to study the interactions. To optimize the experimental conditions that resolution with conditions resembling the native state, wtTel23c was dissolved in different concentrations of cell lysate ranging from 0 mg/mL to 100 mg/mL (Fig. S6), yielding a 75 mg/mL cell lysate as optimal conditions for NMR studies. The stability of wtTel23c in cell lysate was assessed by comparing the spectra of ^15^N-labeled wtTel23c before and after a 12-hour incubation in cell lysate (Fig. S7). Analysis of the 2D ^1^H-^15^N sfHMQC spectra of the imino region revealed that as the cell lysate concentration increased, some cross-peak intensities decreased, while others increased (Fig. S8). This also suggests that the cellular components influence the proportion distribution of conformational states of wtTel23c.

The ex vivo NMR spectra exhibited distinct spectra when compared to samples in dilute solution (Fig. 3A, 3D and 4A). To determine the topology of wtTel23c in the cell lysate, a spectral comparison approach developed by the Volker Dötsch group was used[22]. NMR spectra of telomeric G4 with similar DNA sequences and known structures were collected in dilute solution and compared with the wtTel23c spectrum in cell lysate. The wtTel23c spectrum in cell lysate closely resembled that of wtTel23 in dilute solution, indicating the hybrid-1 conformation (Fig. S9). Further, we used the NMR assignments of wtTel23 in solution to conduct the site-specific assignments of wtTel23c in cell lysate (Fig. 4C).

**Figure 4.**
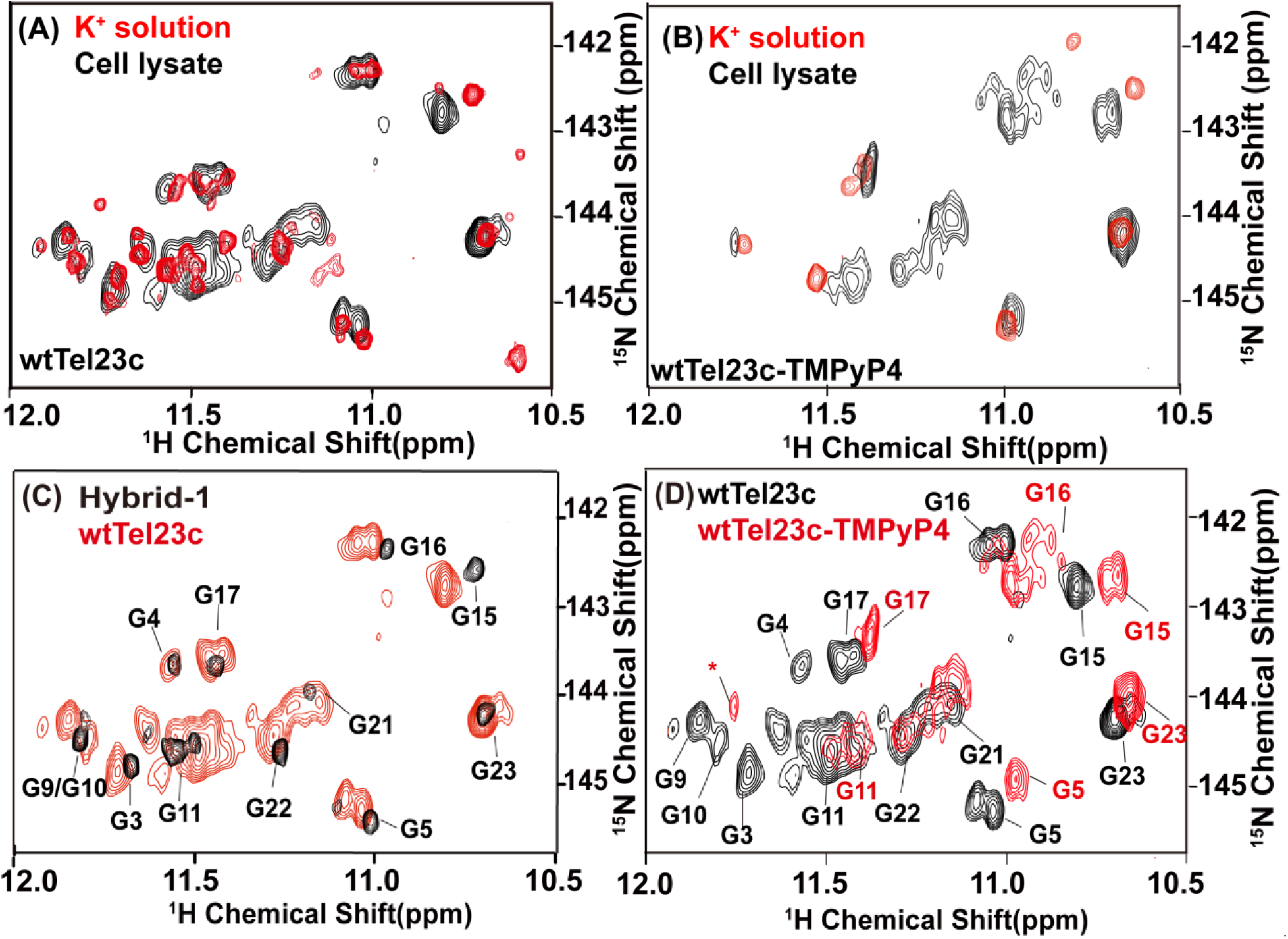
Comparison of ^1^H−^15^N sfHMQC of ^15^N-labeled wtTel23c in different solution conditions. (A) Comparison of ^15^N-labeled wtTel23c in K^+^ buffer (red) and in cell lysate (black). (B) Overlay of ^15^N-labeled wtTel23c- TMPyP4 in K^+^ buffer (red) and in cell lysate (black). (C) Comparison of ^1^H-^15^N sfHMQC of ^15^N-labeled wtTel23c in cell lysate (red) and wtTel23 in solution (black). The assignments of wtTel23c are highlighted on spectrum. (D) Comparison of ^1^H-^15^N sfHMQC spectra of the wtTel23c (black) and the wtTel23c-TMPyP4 (red) in cell lysate. The assignments of wtTel23c and TMPyP4-wtTel23c in cell lysate are labeled in black and red, respectively. The star indicates the unassigned signal.

NMR titration experiments were also conducted to investigate wtTel23c-TMPyP4 interactions in cell lysate. The TMPyP4 was added stepwise into cell lysate solution with ^15^N-labeled wtTel23c to reach 1:0.5 and 1:1.1 molar ratio of ^15^N-wtTel23c and TMPyP4 (Fig. S10). Similar to observation in diluted solution, progress changes in spectral patterns were detected. Addition of TMPyP4 to wtTel23c led reduced number of cross peaks of wtTel23c. The cross-peak with 1H and 15N chemical shifts of 10.9 and 145 ppm, respectively, was used to follow the completeness of the reaction. Without TMPyP4 and with the 1:0.5 molar ratio of TMPyP4 to wtTel23c, two peaks appeared in this position, while at the 1:1.1 molar ratio of TMPyP4 to wtTel23c, a single peak remained, indicative of full binding of wtTel23c with TMPyP4. Further addition of TMPyP4 severely attenuated the intensity of all cross-peaks of wtTel23c, suggesting the presence of multiple binding sites of TMPyP4 on wtTel23c with excess TMPyP4 in cell lysate, which is also similar to that in diluted solution. Thus, we used 1:1.1 molar ratio of TMPyP4 and wtTel23c for further ex vivo NMR studies.

Interestingly, the spectra of wtTel23c-TMPyP4 in cell lysate showed different spectral patterns with the aforementioned wtTel23c-TMPyP4 spectra in diluted solution, hinting different TMPyP4 binding site in cellular environments (Fig. 3B, 3E and 4B). Fig 4D compared the ^1^H-^15^N sfHMQC spectra of unbound wtTel23c and wtTel23c-TMPyP4 in cell lysate. It was found that signals of G3, G4, G9, and G10 were severely weakened or disappearance upon TMPyP4 binding, while G5, G16, G15, and G17 underwent chemical shift changes in either ^15^N or ^1^H dimension (Fig. 4D). In contrast, G21, G22 and G23 showed nearly no changes upon TMPyP4 interactions. Mapped onto the wtTel23 topology (Fig. 3F), G3, G4, G9, and G10 located at the same groove of the G4, adjacent to the propeller loop, thus suggesting that TMPyP4 binds to the nucleotides encompassing the groove. As discussed above, in diluted solution, TMPyP4 binds wtTel23c at the top tetrad. Thus these data clearly demonstrated that the cell lysate affects the binding topology of TMPyP4 at wtTel23c.

Similar to studies in dilute solutions, steady-state fluorescence spectroscopy of wtTel23c with 2-aminopurine mutations was performed in cell lysate to verify the ligand binding region. As shown in Fig 3(H), the fluorescence spectra of A13p and A20p did not change upon binding to TMPyP4, while A2p and A8p from the side composed of G3, G4, G9, and G10 exhibited marked changes after TMPyP4 binding. These results support the findings by NMR, suggesting that TMPyP4 binds to the groove composed of G3, G4, G9, and G10. Considering that in the dilute solution, A2p, A8p, and A20p were affected, but A13p remained unaffected, it showed the discrepancy of the TMPyP4 binding site on wtTel23c between in solution and in cell lysate.

### 3.5. MD simulations to compare the interaction between the wtTel23c and TMPyP4 in cell lysate

To visualize the structures of wtTel23c-TMPyP4 complexes in these two different conditions, we performed MD simulations from an unbound state of a wtTel23c and the ligand (Fig. 5). The simulation system of cell environment is generated based on the unbound configuration of a DNA G-quadruplex and a TMPyP4 ligand, while the simulation system of dilute solution is constructed by adding 150 mM KCl in the simulation box with a DNA G-quadruplex and an unbound ligand. The model of wtTel23c-TMPyP4 in dilute solution showed that an *N*-methylpyridyl group of TMPyP4 places on the top of G9 at the top tetrad, forming π-π stacking with the guanine group of G9. The methyl group is close to the H2 atom of A2 and A20, with a distance of 3.3 Å and 2.5 Å, respectively. The stacking may affect the top tetrad of wtTel23c. Differently, in the model of wtTel23c-TMPyP4 in cell lysate, instead of forming π- π stacking between TMPyP4 and guanine group of tetrad, the TMPyP4 bridges the 5’-end and on the loop of T6-A8 of wtTel23c. Correspondingly, the G3 and the G9 connect the 5’-end and the A8, and spatially close to the TMPyP4 binding sites, which may induce the line-broadening as observed by NMR.

**Figure 5.**
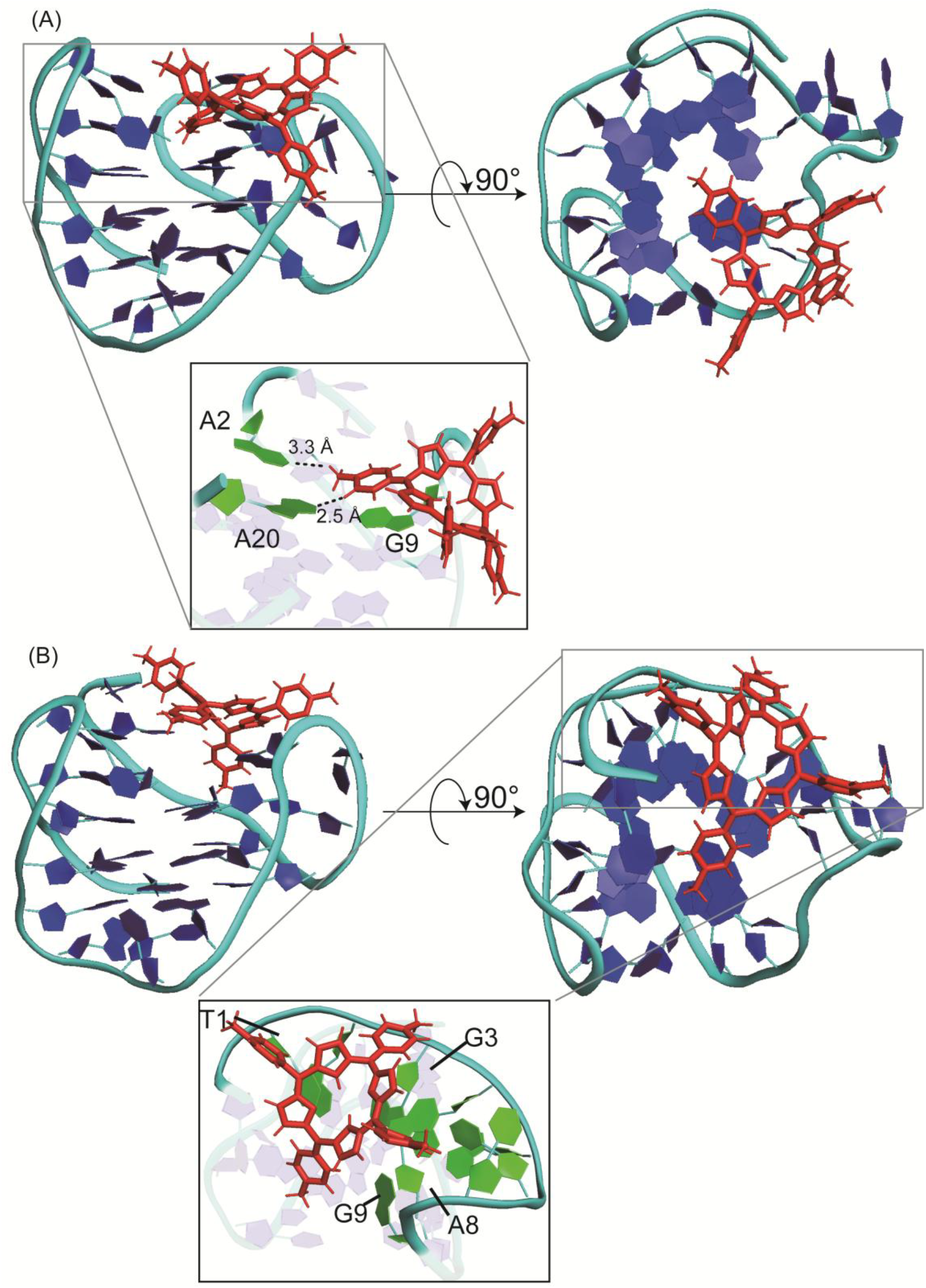
Representing model of wtTel23c-TMPyP4 in dilute solution and in cell lysate environment. (A) Side and front view of wtTel23c-TMPyP4 in dilute solution. (B) Side and front view of wtTel23c-TMPyP4 in cell lysate. The TMPyP4 is shown in red. The wtTel23c are shown in blue and cyan. The zoom regions highlight the contacts between TMPyP4 and wtTel23c.

We calculated the root-mean-square deviation (RMSD) of heavy atoms for both wtTel23c and ligand to examine whether the simulations reach a stable state and the bound state remain in the steady form. As shown in Fig 6, the simulations in the two conditions were both convergent after a 500 ns production run. The stable bound states of TMPyP4 on wtTel23c in the two conditions were then extracted from our trajectories and illustrated in Fig. 6A and Fig. 6E, respectively, where the ligand binds to the propeller loop (G3, G4, G5, T6, T7, A8, G9, G10, and G11) of G4 in the cell lysate environment but prefers to the top stacking (G3, G9, G17, and G21) binding mode in the dilute solution. To characterize the binding sites of TMPyP4 on wtTel23c in the two different conditions, the number of hydrogen bonds (N1-H1…O6) in the three G-tetrads (top layer G3-G9-G17-G21, middle layer G4-G10-G16-G22, and bottom layer G5-G11-G15-G23) were plotted against simulation time (Fig. 6C and 6G). In the model simulating the wtTel23c-TMPyP4 in cell lysate environment, the number of hydrogen bonds remained stable in the top and bottom layers, while a decrease in the number of hydrogen bonds was observed in the middle layer. This disruption might be induced by the interaction between TMPyP4 on wtTel23c, where the binding site is close to G4, G10 positions of the middle layer. In the dilute solution, the number of hydrogen bonds was decreased in the top layer, demonstrating that TMPyP4 mainly binds to the top layer of wtTel23c. We also calculated the Root-mean-square fluctuation (RMSF) for each DNA residue to evaluate the effects for the binding of TMPyP4 on wtTel23c. RMSF characterizes the local structural fluctuations before and after the binding of ligand. As shown in Fig. 6D and Fig. 6H, there are three peaks (T6-A8, T12-A14, and T18-A20) for DNA in the two conditions. In the cell lysate environment, the binding of ligand slightly increased the RMSF values of first two peaks but reduced the third peak and the first data point. However, in the dilute solution, the binding of ligand significantly reduced the RMSF values of all three peaks. These results indicate that TMPyP4 might stabilize the complex structure in the dilute solution.

**Figure 6.**
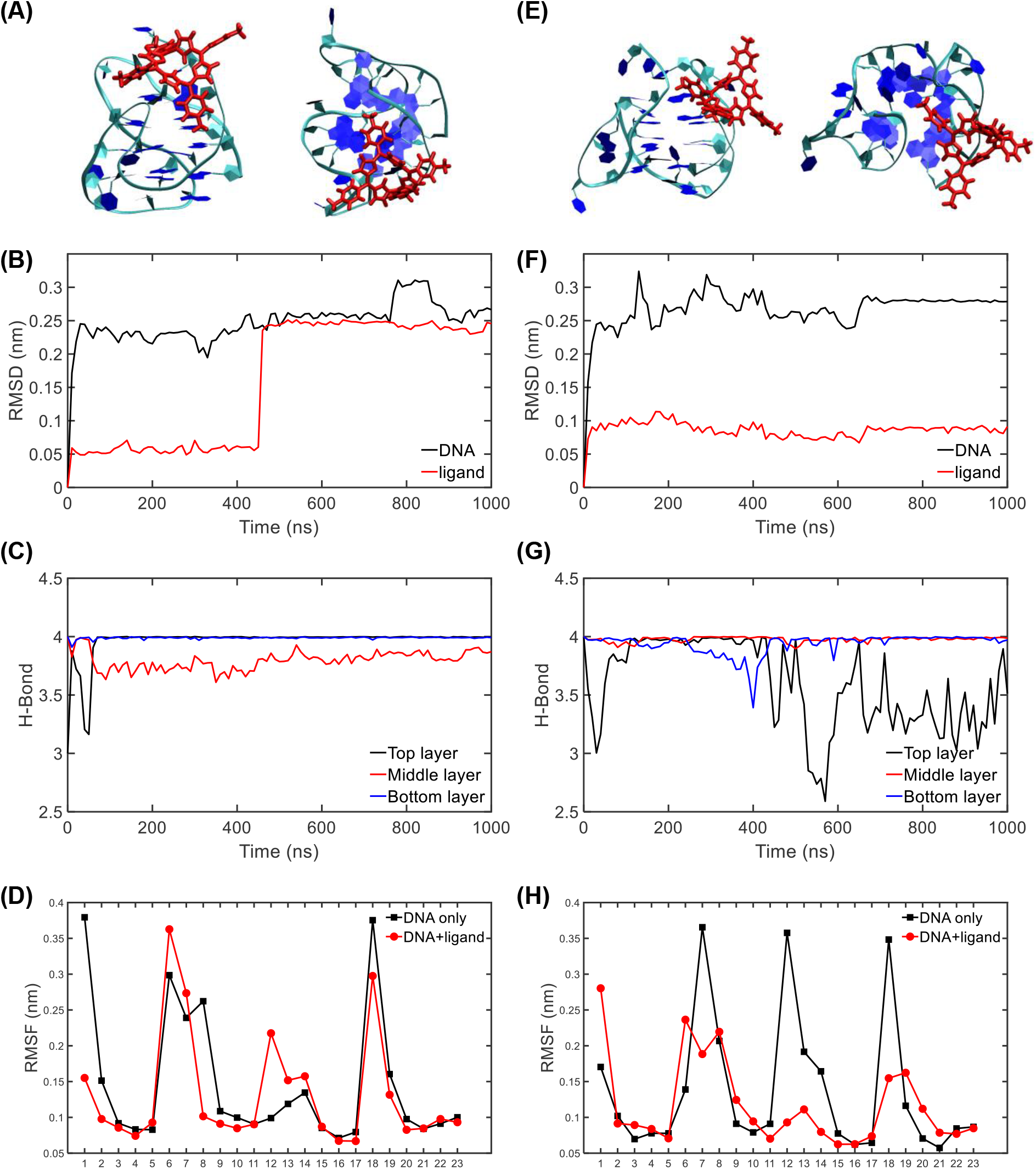
Snapshots and the order parameters for wtTel23c-TMPyP4 in the cell lysate environment and the dilute solution from simulations. (A) Front view (left) and top view (right) of the binding structure, (B) RMSD of the DNA/ligand with respect to the starting structure, (C) number of hydrogen bonds (N1-H1…O6) in the three G-tetrad layers, (D) RMSF of each DNA residue for wtTel23c-TMPyP4 in the cell lysate environment. (E-H) Similar results for wtTel23c-TMPyP4 except in the dilute solution.

### 3.6. Evaluation of the cellular components affecting the G4 and TMPyP4 binding

The binding site of wtTel23c-TMPyP4 varies between solution and cell lysate, suggesting potential influences from metabolic compounds or interactions with cellular proteins. To investigate the factors affecting structural polymorphism and TMPyP4 binding in cellular environments, we treated the occyte exctracts to obtain two different samples, including cleared cell lysates and crowded lysates. The cleared cell lysates were prepared by heating the lysate suspend solution and removing thermal denatured proteins, in which most of the proteins are removed. In this condition, the metabolic compounds, nucleic acids and the thermal stable proteins are preserved. Thus one could evaluate how these components influence on the wtTel23c topology and its interactions with ligands. The crowded lysate environment was created by adding of 2 mM NA-wtTel23c to the cell extract, acting as a “bait” to capture all potential binding partners from the extract.[22] In this condition, one could evaluate exclusively the effects of crowdness on wtTel23c structure and interactions.

We conducted three sets of experiments on wtTel23c in the presence of cellular components. Firstly, we dissolved and collected the sfHMQC spectra of ^15^N-labeled wtTel23c in cleared cell lysate, resulting in a spectrum similar to wtTel23c in the cell lysate, but distinct from the that in dilute solution (Fig. S11). This indicates that metabolic cellular compounds that are stable in thermal denaturation has a similar effect on the conformation of wtTel23c as observed in cell lysate, demonstrating that those components affect the conformation polymorphisms of ligand free wtTel23c. Secondly, we collected the sfHMQC spectra of ^15^N-labeled wtTel23c in the crowded lysates. It closely resembles those in the cell lysate (Fig S11C and S11D), indicating that the crowded environment induces similar effects on the topology of the wtTel23c as the cell lysate does. These findings suggest that specific components in cell lysate resistant to thermal denaturation are critical in affecting the conformations of ligand free wtTel23c.

Next, we performed TMPyP4 titration experiments in the cleared lysate and crowded lysate environment (Fig. 7). Interestingly, both of them exhibited a similar pattern to that observed in the dilute solution (Fig. 7A), but distinctly different from those in the cell lysate environment (Fig. 7D). Both cleared and crowded lysate states essentially block binding between the cellular components and wtTel23c or removed the most proteins that may binds wtTel23c. This suggests that the proteins precipitated after thermal denaturation in cell extracts may influence TMPyP4 binding to wtTel23c, rather than the crowdedness environment.

**Figure 7.**
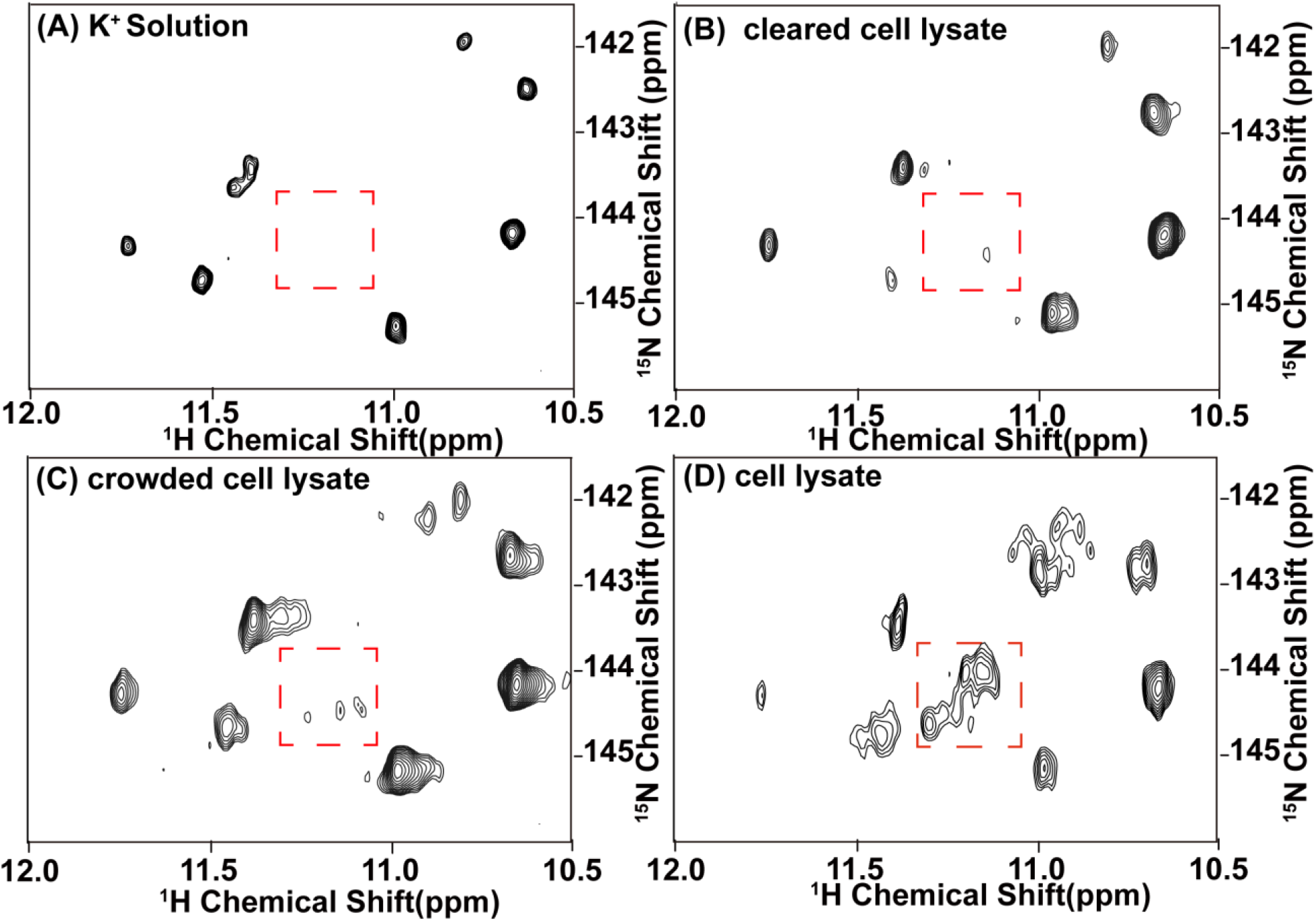
Comparison the interaction of TMPyP4 and wtTel23c under different conditions. 2D ^1^H-^15^N sfHMQC of wtTel23c-TMPyP4 in K^+^ dilute solution (A), in cleared cell lysate (B), in crowded cell lysate (C) and in cell lysate environment (D) The box indicates significantly different signals in different condition.

To further confirm this conclusion, we conducted the third set of experiments (Fig. 8A). TMPyP4 and wtTel23c were initially dissolved in the cleared cell lysate (Fig. 8B), showing the spectral close that in the dilute solution (Fig. 8C). Further the cell lysate powder was added into this solution to reach 75 mg/mL protein concentrations. We obtained a spectral signal similar to that of wtTel23c-TMPyP4 supplemented in cell lysate directly. (Fig. 8D). The replenished extract altered the spectral pattern of wtTel23c-TMPyP4 from a dilute solution-like state to a pattern resembling the cell lysate state. Thus the supplement of cell extract powders recover the spectral patterns of wtTel23c-TMPyP4 in cellular states. Taken together, these observations indicate that the interactions between cell lystate proteins and wtTel23c, rather than the crowdedness environment can modulate the ligand-binding site on wtTel23c.

**Figure 8.**
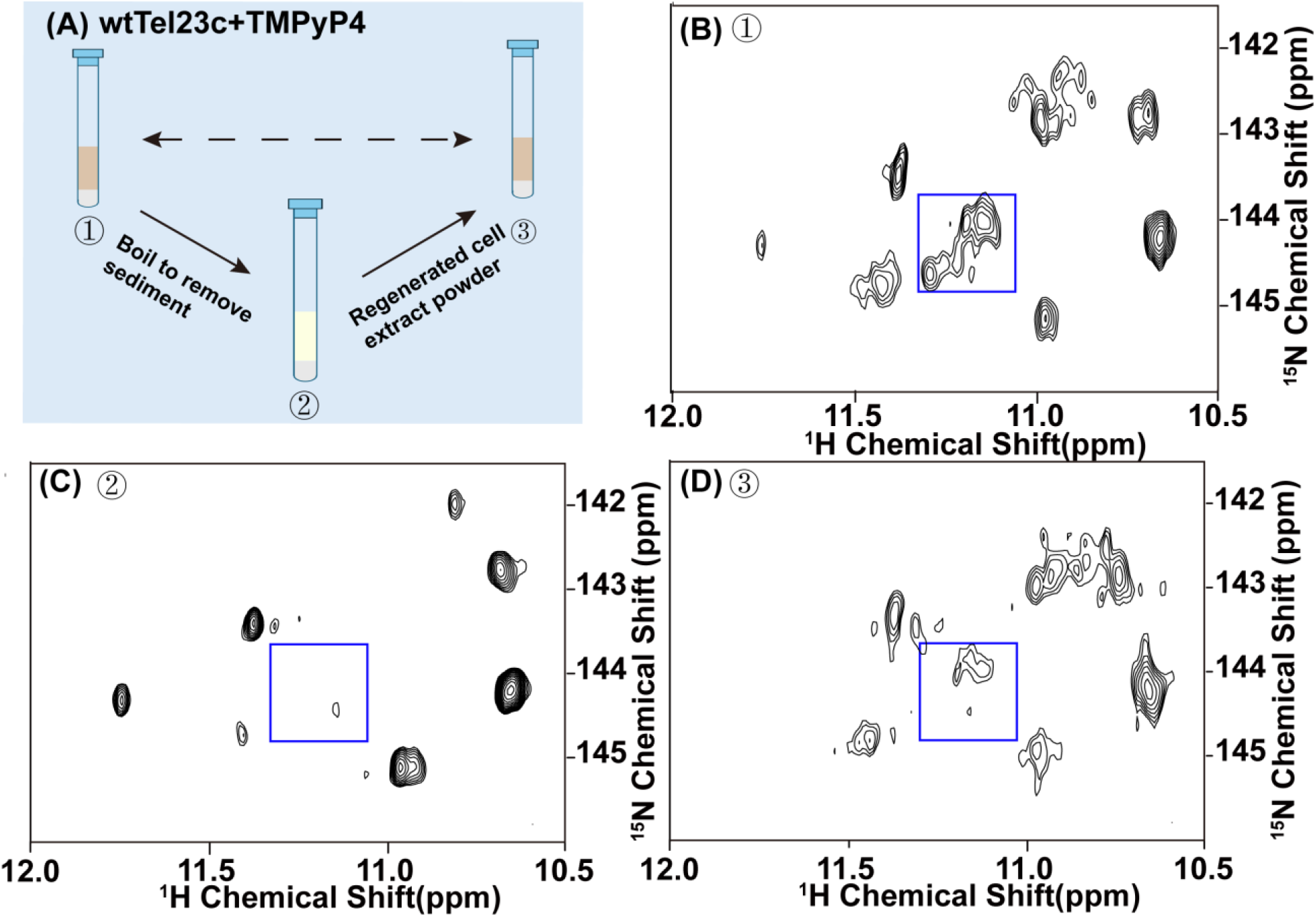
Cell lysate effect on the binding site of wtTel23c-TMPyP4. (A) The steps of continuous reaction of wtTel23c-TMPyP4.Step ①, wtTel23c interact with TMPyP4 in 75 mg/mL cell lysate, and corresponding spectrum. (B) Step ②, boiling sample 1 to remove sediment, and corresponding spectrum. (C) Step ③, add cell extract to sample ② until the final concentration is 75 mg/mL, and corresponding spectrum (D) The box indicates significantly different signals in different condition.

## 4. Conclusions

In this study, we present a novel biosynthesis method for ^15^N labeling of ssDNA, which combined with enzymatic digestion to obtain the targe products. This method has several lines of advantages. First, it is suitable for the synthesis of ssDNA even with high GC content or repeat units. Second, this approach is economical compared to chemical synthesis or enzymatic methods, as it uses the low cost ^15^N-NH_4_Cl as ^15^N-labeling metabolic precursor. Third, this method achieves uniformly ^15^N-labeling, and yielded sub-milligram quantities of ^15^N-labeled DNA in each liter of cell culture, close to the level of ^15^N-labeled protein expression by *E. coli*. These merits satisfied the requirements for NMR studies and allow the interaction studies on DNA and ligands in vivo or in the cell lysate environment.

The method was used to study the topology of telomeric G4 and its complex with ligands. Telomeric G4 has been known to adopt different topologies depending on solution conditions, sequences, and as well the presence of the cellular context. It has also been debated about the ligand binding sites of telomeric G4. Here, using the 15N-labeled wtTel23c showed that cellular environment has significant impact on the conformation equilibrium of telomeric G4, and interestingly, the binding sites of TMPyP4 ligand to the telomeric G4.

Evaluation on the factors that the binding model of wtTel23c-TMPyP4 suggest that possible protein-wtTel23c interactions may determine the binding sites between wtTel23c and TMPyP4. According to the previous reports, telomeric DNA is associated with shelterin proteins including TRF1, TRF2, RAP1, TIN2, TPP1 and POT1 which protect the chromosomal ends via physical association[42-44]. Especially, Katrin Paeschke’s group has indicated that both TEBPα and TEBPβ are required for G4 formation in vivo[45]. Sujay Ray and his team demonstrated that POT1/TPP1 interaction with G4 plays a protective role in safeguarding telomeric G4 structures from binding by replication protein A, thereby preventing the generation of DNA damage signals[46]. Other G4 binding proteins may also binds to telomeric G4 and consequently affects the binding mode of telomeric G4 and ligands. It thus may raise a challenging in fragment based drug-design using the structures of G4-ligand complexes without the protein partners. It is also intriguing to develop methods of G4-ligand screening in cellular conditions, in which the interactions between G4 and proteins are intrinsically considered.

In addition, this work provides a practical approach to prepare ^15^N ssDNA for in vitro and in vivo studies. One can extend this approach to produce ^13^C-labeled ssDNA or sparse 13C-labeled ssDNA as many protein NMR studies have developed, using ^13^C-glucose or other ^13^C-labeled carbon source. The research scope of this work can also be expanded to other types of DNAs, i.e, i-motif, to unrevealing the structure, functions and mechanisms of biological processes.

## Supporting information

This PDF file includes: Figs. S1 to S11 Tables S1

## Conflict of interest

The authors declare that they have no conflict of interest.

### Acknowledgments

All of the NMR experiments were conducted at the Beijing NMR Center, the NMR Facility at the National Center for Protein Sciences at Peking University, the National Center for Protein Science (Shanghai), and the 800 MHz spectrometer at East China University of Science and Technology. The work was supported by the National Key R&D Program of China (2024YFA0917100), the National Natural Science Foundation of China (22274050, 21925406), the Shanghai Science and Technology Commission (contract number: 23J21900300, 24HC2810700) and the Fundamental Research Funds for the Central Universities. We are grateful for the assistance of Dr. Xiaogang Niu and Dr. Hongwei Li of Peking University and Dr. Zhi-jun Liu of the National Center for Protein Science, Shanghai, on NMR data collection. We also thank the staff members of the Nuclear Magnetic Resonance System at the National Facility for Protein Science in Shanghai (NFPS), Shanghai Advanced Research Institute, Chinese Academy of Sciences, China for providing technical support and assistance in data collection and analysis.

## Author contributions

M. Z. was primarily responsible for the experimental design, experimental validation, and manuscript writing. F. T., S. Z. and L. D. were mainly responsible for the MD simulations. M. Z., C. M. and M.J. produced DNA samples. W. Z. and X. L. provided the oocyte powder samples. M. Z. provided conceptions of the research and guided the experimental design. C. L. and S. W were responsible for designing the overall research approach and provided constructive guidance.

