## Supplementary material for "Bio-synthesis of ^15^N-Labeled G-Quadruplexes to Investigate the Structure and Interactions in the Cell Lysate using Nuclear Magnetic Resonance": This PDF file includes: Figs. S1 to S11 Tables S1

Target sequence unit 1

GGTACCTAGGGTTAGGGTTAGGGTTAGGGGTACCGGTACCTAGGGTTAGGGTTAGGGTTAGGGGTACCGGTACCTAG GGTTAGGGTTAGGGTTAGGGGTACCGGTACC

Linker 1

Cgccctgctttctcggcatctctgatagcctgagaagaaaccccaactaaatccgctgcttcacctattctccagcgccgggttattttcctcgcttccgggctgtcatcattaaactgtgcaatgg cgatagccttcgtcatttcatgaccagcgtttatgcactggttaagtgtttccatgagtttcattctgaacat

Target sequence unit 2

GGTACCTAGGGTTAGGGTTAGGGTTAGGGGTACCGGTACCTAGGGTTAGGGTTAGGGTTAGGGGTACCGGTACCTAG GGTTAGGGTTAGGGTTAGGGGTACCGGTACC

Linker 2

gatcggtgggcagtttaccttcatcaaatttgcccattaactcagtttcaatacggtgcagagccagacaggaaggaataatgtcaagccccggccagcaagtgggctttattgcataagtgaca tcgtccttttccccaagatagaaaggcaggagagtgtcttctgcatgaatatgaagatctGGTACCcatccgtga

Target sequence unit 3

GGTACCTAGGGTTAGGGTTAGGGTTAGGGGTACCGGTACCTAGGGTTAGGGTTAGGGTTAGGGGTACCGGTACCTAG GGTTAGGGTTAGGGTTAGGGGTACCGGTACC

Linker 3

Ataatcagaccgacgatacgagtgggaccgtggtcccagtctgattatcagaccgacgatacgagtgggaccgtggtcccagactaataatcagaccgacgatacgagtgggaccgtggt cccagactaataatcagaccgacgatacgagtgggaccgtggtcccagtctgattatcagaccgacgatacaagtggaac

Target sequence unit 4

GGTACCTAGGGTTAGGGTTAGGGTTAGGGGTACCGGTACCTAGGGTTAGGGTTAGGGTTAGGGGTACCGGTACCTAG GGTTAGGGTTAGGGTTAGGGGTACCGGTACC

Linker 3

ttaatacgatacctgcgtcataattgattatttgacgtggtttgatggcctccacgcacgttgtgatatgtagatgataatcattatcactttacgggtcctttccggtgatccgacaggttacggggc ggcgacctcgcgggttttcgctatttatgaaaattttccggtttaaggc

gtttccgttcttcttcgtcata

Target sequence unit 5

GGTACCTAGGGTTAGGGTTAGGGTTAGGGGTACCGGTACCTAGGGTTAGGGTTAGGGTTAGGGGTACCGGTACCTAG GGTTAGGGTTAGGGTTAGGGGTACCGGTACC

Table 1. An illustration of the ligation sequence into PUC57. The underlined sequence represents the target sequence. Kpn I and BamH I restriction enzyme digestion sites are in red and blue respectively. The rest are linker sequences.

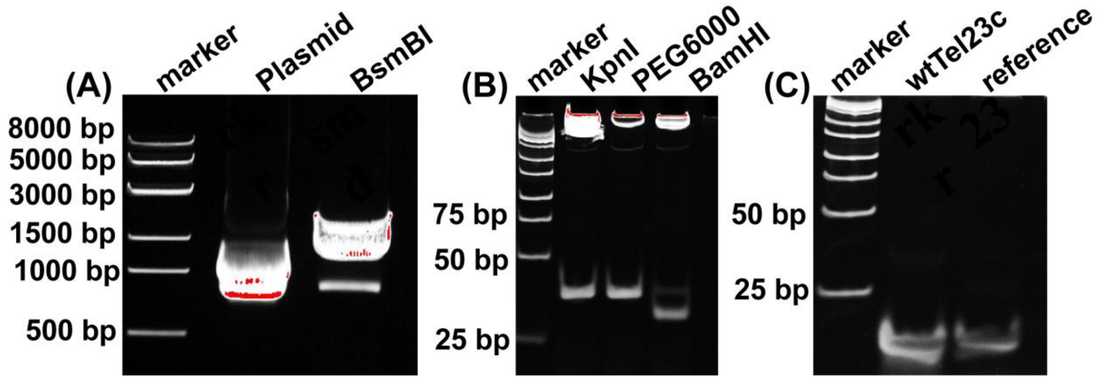


**Figure S1. Characterization of wtTel23c prepared by RED-ssDNA method**. (A) Agarose gel (1%) analysis of the *BsmBI*-digested plasmid harboring the wtTel23c sequences. Aliquots of plasmid with and without *BsmBI* digestion were analyzed on agarose gels followed by gel red staining. (B) Native PAGE (12%) of Kpn I and BamH I-digested plasmid. The plasmid was digested by Kpn I to produce the fragment with the target DNA and the vector fragment (lane Kpn I). The vector fragment was removed using polyethylene glycol 6000, and the target fragment was further digested by BamH I to obtain the target dsDNA (lane BamH I). (C) Native PAGE (12%) analysis of wtTel23c compared with the reference.

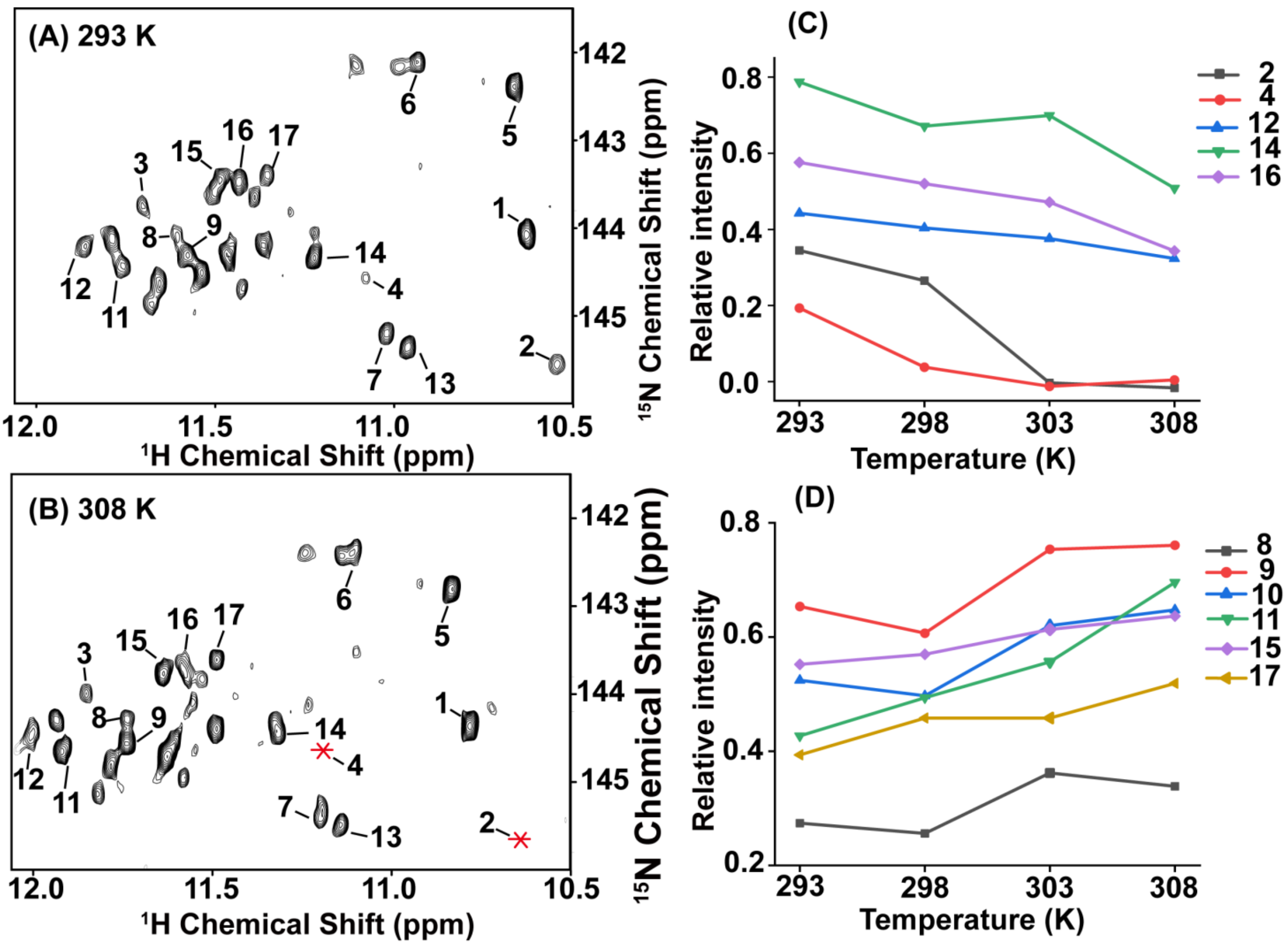


**Figure S2. Temperature dependent conformational equilibrium of $^{15}$N-wtTel23c in dilute solution.** (A,B) $^{1}$H−$^{15}$N sfHMQC spectra of the imino region of wtTel23c at 293K (A) and 308K (B). The numbers highlighted on the spectra are only the numeric numbers of the corresponding peaks (not the sequential assignments). (C,D) Temperature dependent peak intensities. All peak intensities were normalized using the peak intensity of peak1. The asterisk represents the signals disappeared at 308K. (C) The cross-peaks with decreased relative intensities over temperature increasing. (D) he cross-peaks with increased relative intensities over temperature increasing.

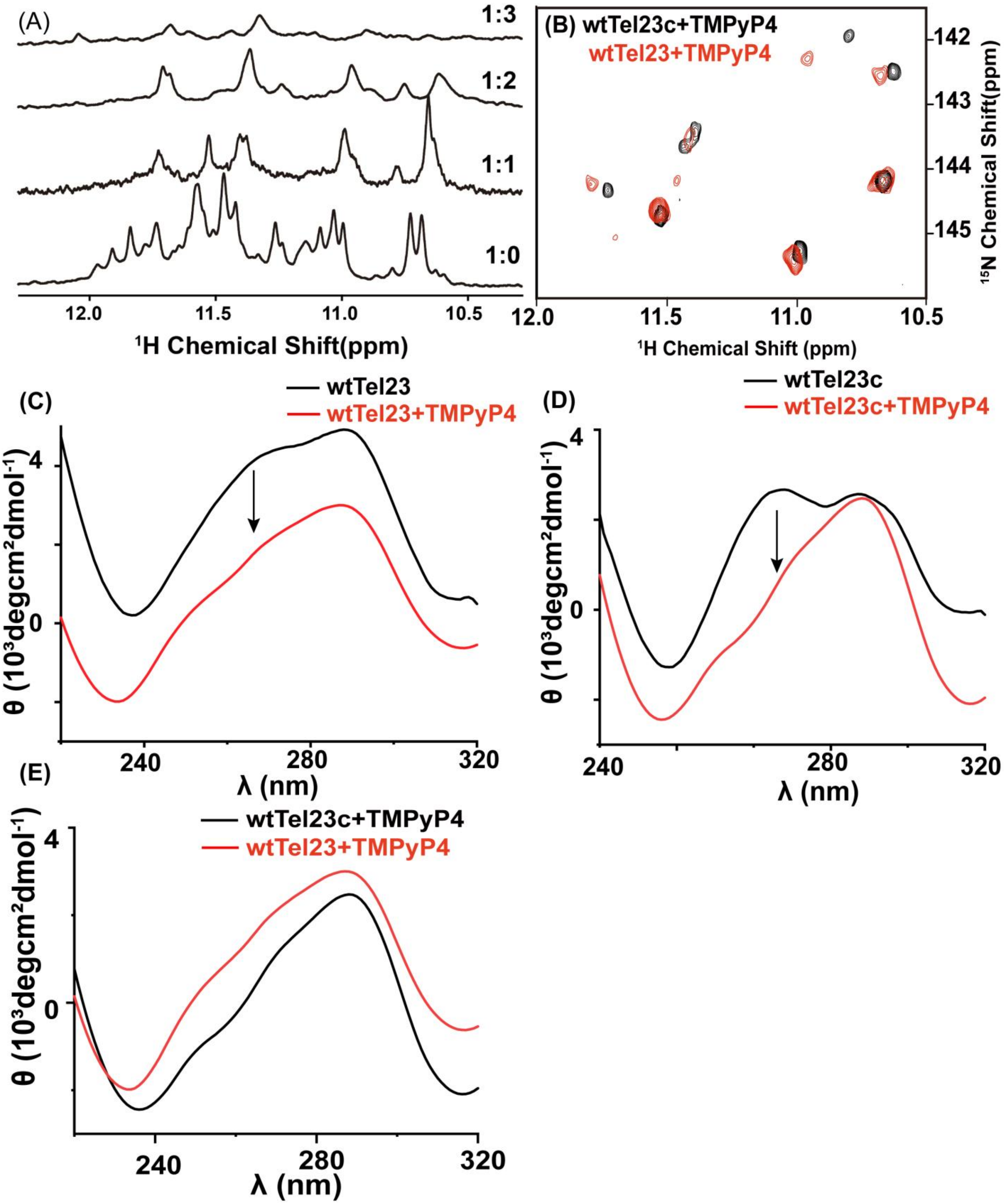


**Figure S3. CD and NMR spectra of wtTel23, wtTel23c and their complex with TMPyP4 ligands.** (A) Titration of TMPyP4 to a molar ratio of wtTel23c vs. TMPyP4 at 1:1, 1:2, and 1:3. (B) Comparison of 2D $^{1}H$-$^{15}N$ sfHMQC NMR spectra with imino regions of wtTel23-TMPyP4 (red) and wtTel23c-TMPyP4 (black) . The signal assignments of wtTel23c-TMPyP4 complex are shown .(C) CD spectra of ligand free wtTel23 (black) and wtTel23-TMPyP4 complex (red). (D) CD spectra of ligand free wtTel23c (black) and wtTel23c-TMPyP4 complex (red).(E) A comparison of CD spectra of wtTel23-TMPyP4 (red) and wtTel23c-TMPyP4 (black).

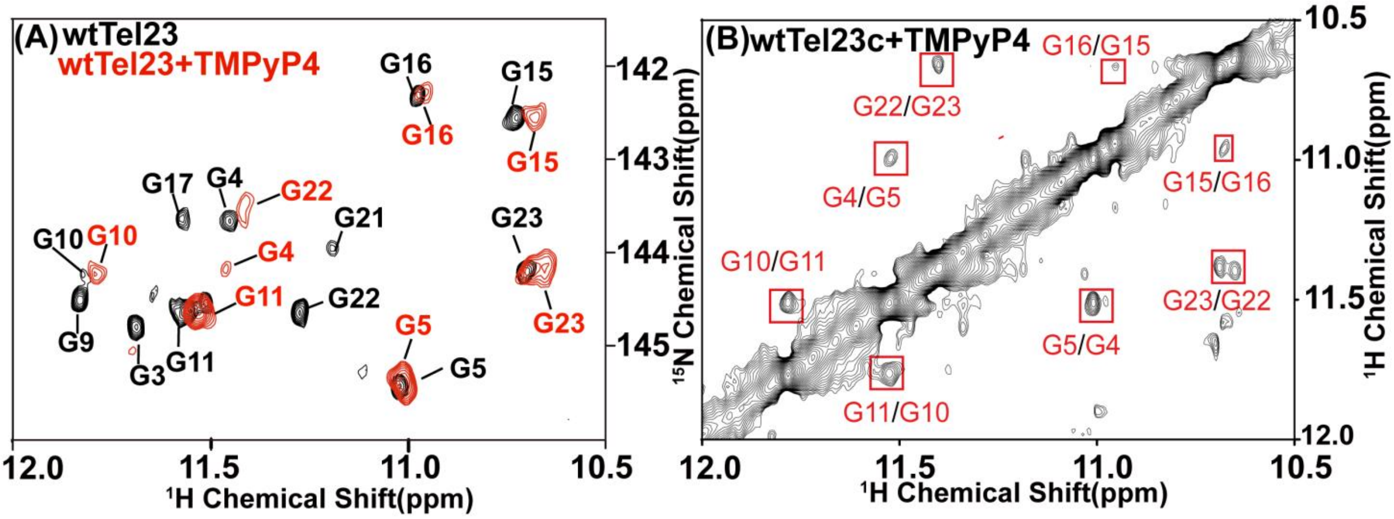


**Figure S4. Assignment of wtTel23-TMTyP4 in solution.** (A) The $^{1}$H-$^{15}$N sfHMQC signal with assignments of ligand free wtTel23 (black) and wtTel23-TMTyP4 (red). The assignments of G16, G15, G23, G5 and G11, were obtained through by spectral comparison. The assignments of other nucleotides were further confirmed by the by $^{1}$H-$^{1}$H NOESY spectrum (B). The $^{1}$H-$^{1}$H NOESY spectrum with mixing time of 200 ms of wtTel23c-TMPyP4 complex, showing $^{1}$H-$^{1}$H connections. Cross-peaks that corresponding to the imino protonts of sequential connected guanines are framed and labeled with the residue numbers.

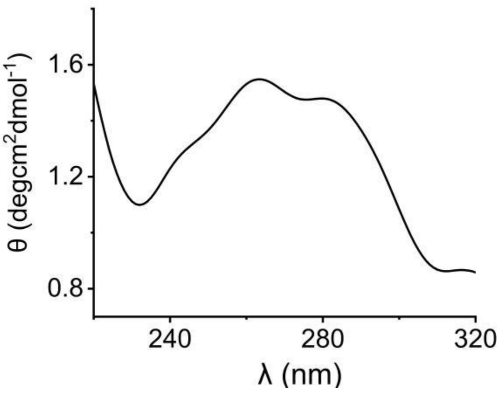


**Figure S5. CD spectra of T13A-wtTel23c**. Spectrum of T13A-wtTel23c with positive bands around 285 and 260 nm, and negative bands around 240 nm, indicating of a hybrid-1 topology.

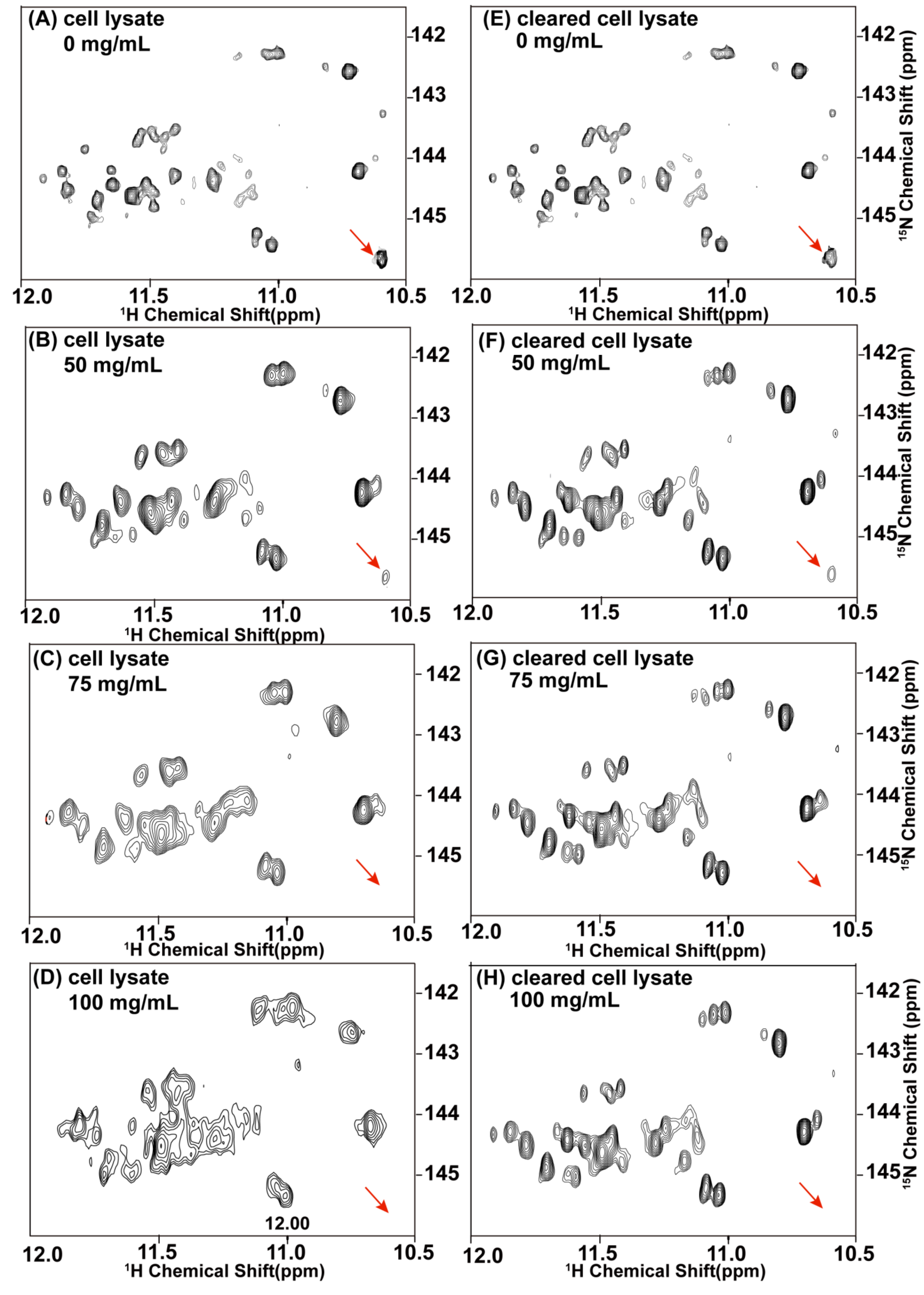


**Figure S6. The influence of cell lysate and cleared lysate on the conformations of wtTel23c.** (A-D) sfHMQC spectra of $^{15}$NwtTel23c in different concentrations of cell lysate. The concentration of cell lysate are shown on the spectra. (E-H) sfHMQC of wtTel23c in cleared lysate. The red arrows indicate a signal weakened and disappeared upon increasing of cell lysate. All spectra were acquired at 298k.

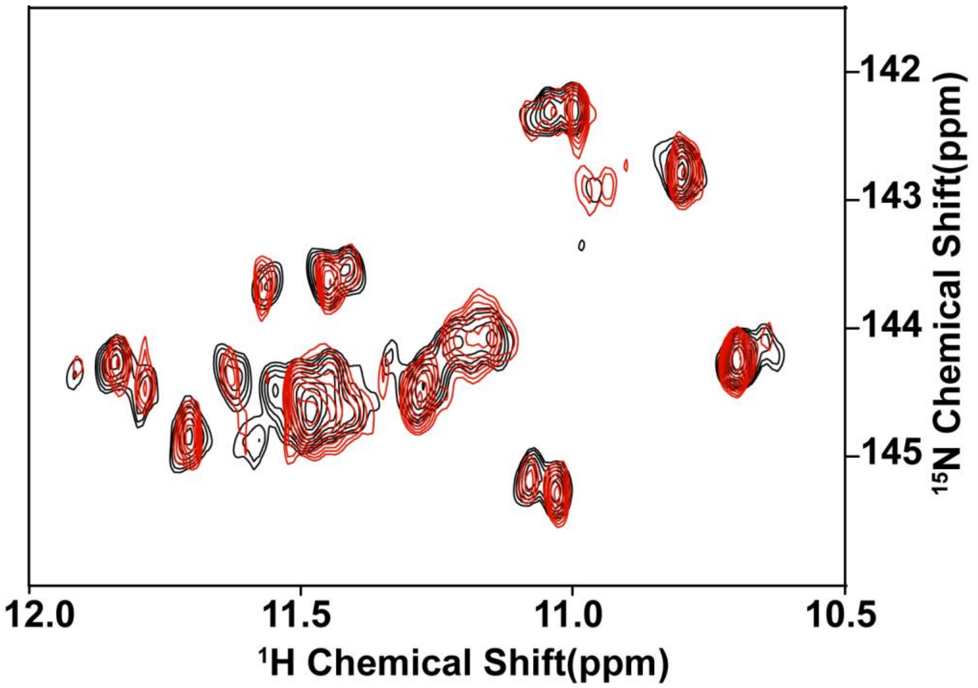


**Figure S7. Stability characterization of wtTel23c in cell lysate.** The sfHMQC spectra $^{15}$N-labeled wtTel23c collected immediately dissolved in cell lysate (black) and after 12 h incubation (red).

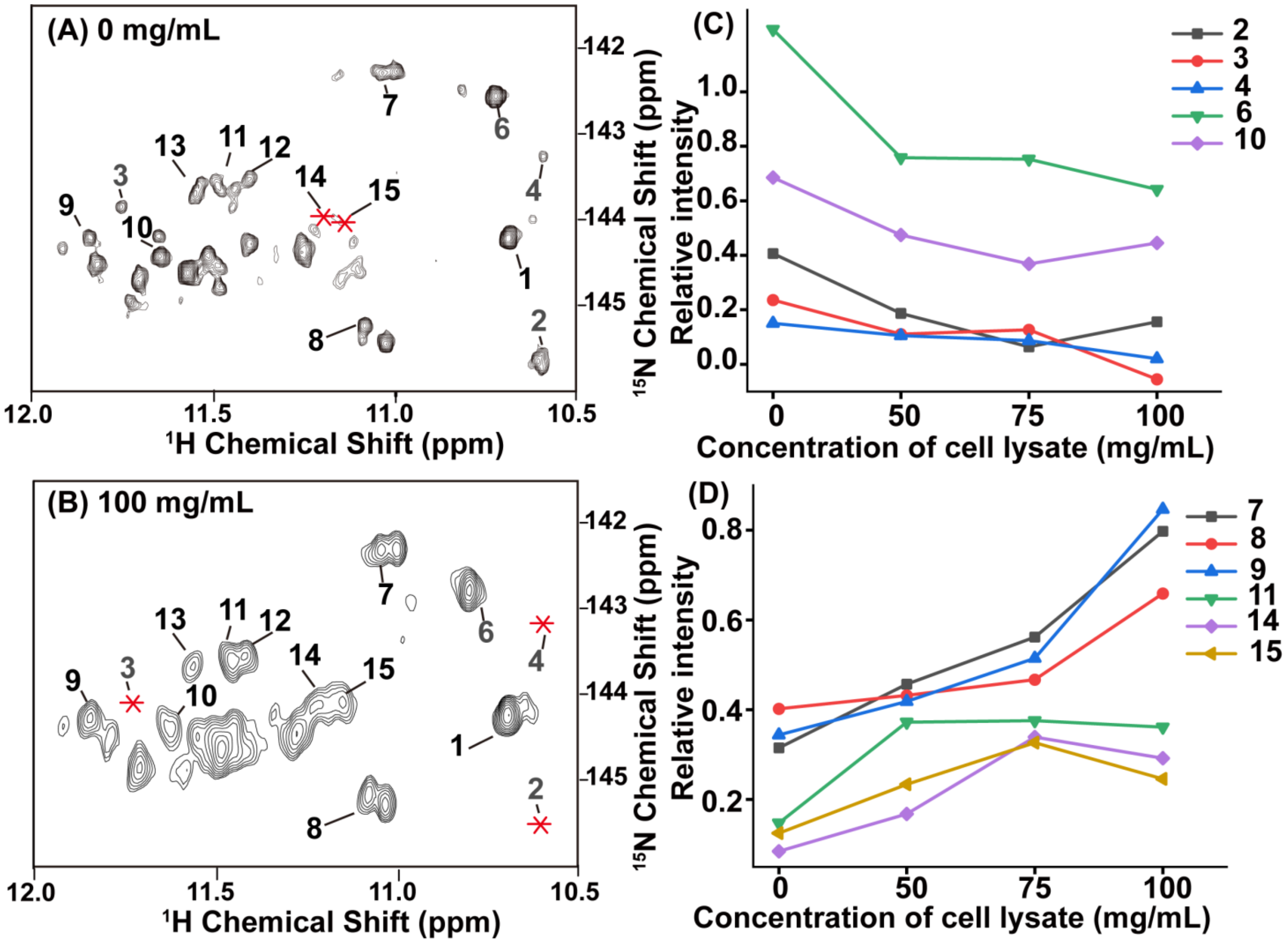


**Figure S8. Analyzing the effect of cell lysate concentration on wtTel23c conformation by 2D $^{1}$H–$^{15}$N sfHMQC spectra of the imino region.** (A) $^{1}$H-$^{15}$N sfHMQC spectrum of wtTel23c in dilute solution. (B) $^{1}$H-$^{15}$N sfHMQC spectrum of wttel23c at 75 mg/ml cell lysate. With peak 1 as the reference, the corresponding intensities of other signal peaks relative to peak 1 are calculated respectively. The label of the peak in the spectrum is a random number, which is independent of the DNA primary sequence number to which the NMR signal belongs. The asterisk represents the disappeared signal. The change trend curve of the relative intensity of the corresponding number of signals under different concentration of cell lysate, (C) represents the signal peak that the relative intensity of the signal increases with the increase of the cell lysate concentration; (D) A signal peak with a decrease in the relative strength of the signal.

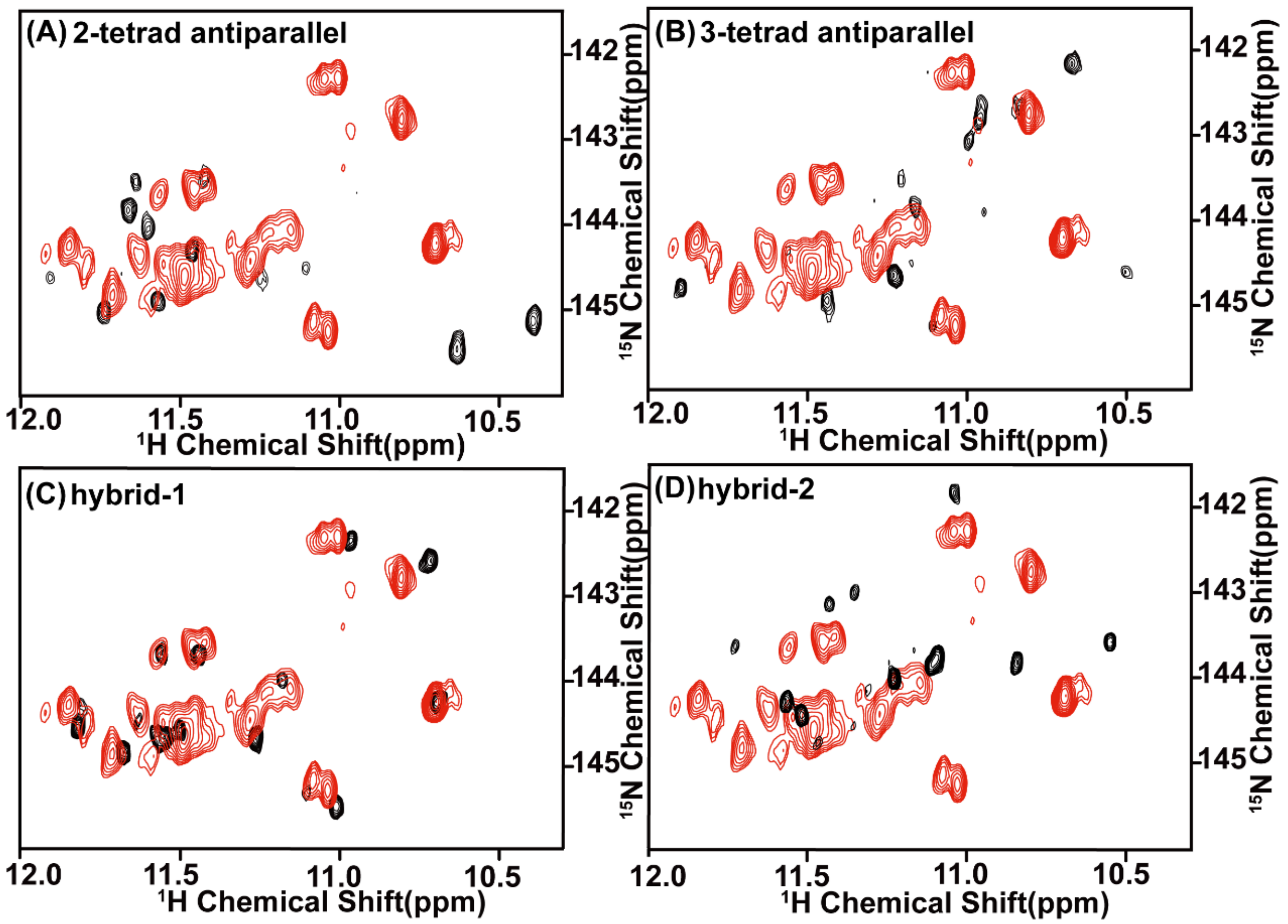


**Figure S9. Comparison of wtTel23c in cell lysates and other telomeric G4 with known structures under dilute solution.** The spectra of wtTel23c in cell lysate are shown in red. (A-D) The spectra of other telomeric G4 with known spectra are shown in black, including and reported G4s topology in dilute buffer, (A) d(G3(TTAGGG)3T) sequence fold into 2-tetrad antiparallel topology in dilute $K^+$-contain buffer, (B) d(AG3(TTAGGG)3) sequence folded into 3-tetrad antiparallel topology in dilute $Na^+$-contain solution, (C) d(AG3(TTAGGG)3) folded into hybrid-1 topology in dilute $K^+$-containing buffer, (D) ) d(TTAG3(TTAGGG)3TT) sequence fold hybrid-2 topology in dilute $K^+$-containing buffer.

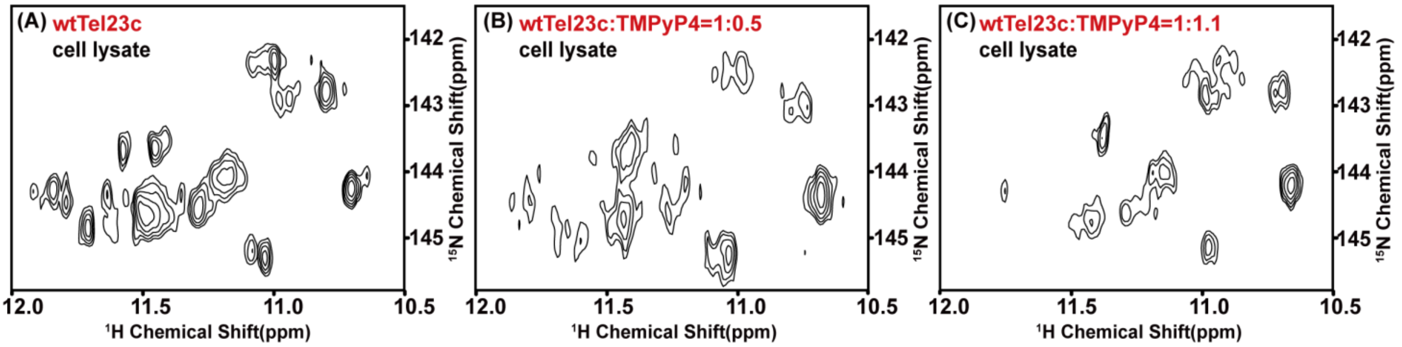


**Figure S10. NMR Titration of $^{15}$N-wtTel23c and TMPyP4.** (A) 2D $^{1}$H-$^{15}$N sfHMQC of the wtTel23c in lysate without TMPyP4 (A). (B-C) 2D $^{1}$H-$^{15}$N sfHMQC of the wtTel23c in lysate with TMPyP4 in a ratio of wtTel23c:TMPyP4 at 1:0.5 (B) and 1:1 (C), respectively.

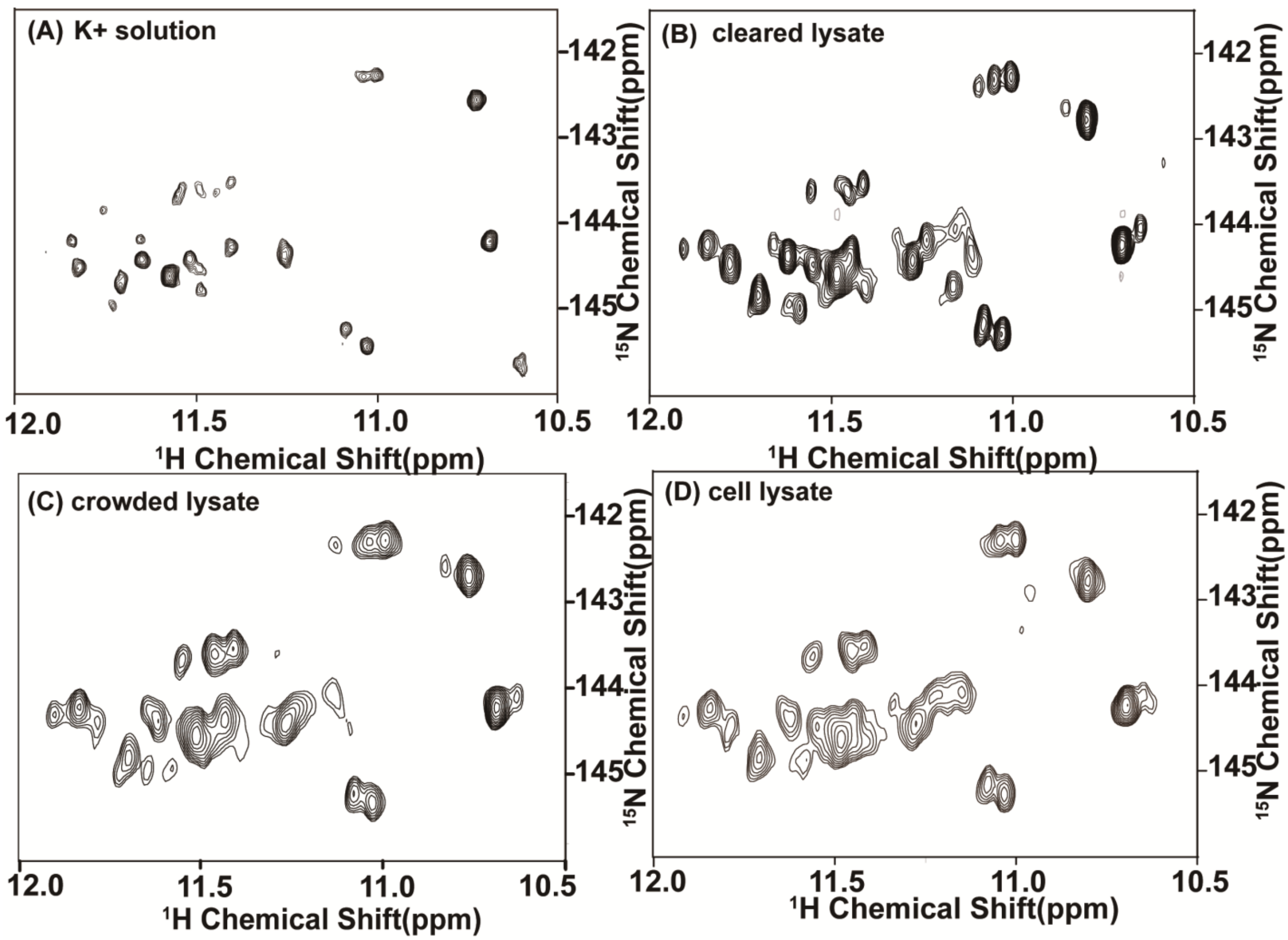


**Figure S11. 2D $^{1}$H-$^{15}$N sfHMQC of the wtTel23c in different conditions.**
